# The influence of heavy metal stress on the evolutionary transition of teosinte to maize

**DOI:** 10.1101/2025.03.17.643647

**Authors:** Jonathan Acosta-Bayona, Miguel Vallebueno-Estrada, Jean-Philippe Vielle-Calzada

## Abstract

Maize originated from teosinte *parviglumis* following a subspeciation event occurred in volcanic regions of Mesoamerica. The elucidation of the phenotypic changes that gave rise to maize have focused on the direct consequences of domestication, with no insights on how environmental factors could have influenced specific gene function and human selection. The genome of the *Palomero toluqueño* landrace suggested heavy metal (HM) effects on maize domestication (Vielle-Calzada et al., 2009). To test the hypothesis that HM stress influenced the evolutionary transition of teosinte to maize, we exposed both subspecies to sublethal concentrations of copper and cadmium. We also assessed the genetic diversity of three HM response genes mapping to chromosome 5 (chr.5) and previously shown to be affected by domestication: *ZmHMA1*, *ZmHMA7* – encoding for heavy metal ATPases of the P1_b_ family-, and *ZmSKUs5,* encoding for a multicopper oxidase. *ZmHMA1* and *ZmSKUs5* are within a genomic region containing hundreds of genes linked to QTLs with pleiotropic effects on domestication. The genomic analysis of chr.5 shows that the three genes were under strong positive selection as compared to previously identified domestication genes. Large-scale transcriptomic comparisons indicate that many other loci containing HM response genes were also positively selected across the maize genome. Exposure of teosinte *parviglumis* to HM stress results in a plant architecture reminiscent of extant maize, and upregulation of *Teosinte branched1* (*Tb1*) in the meristem. *ZmHMA1* and *ZmHMA7* are expressed throughout development and respond to HM stress in both subspecies. *ZmHMA1* is mainly involved in restricting plant height and optimizing the number of female inflorescences and seminal roots. Paleoenvironmental studies reveal a temporal and geographical convergence between volcanic eruptions and maize emergence during the early Holocene, at periods of climatic instability. Overall, these results suggest that abiotic stress influenced the evolutionary transition that gave rise to extant maize through the activity of heavy metal response genes and their phenotypic effects, as a possible consequence of volcanic activity.

## Introduction

Driven by either conscious or unconscious human selection, domestication can be considered a gradual evolutionary process resulting in the adaptation to agroecological environments and anthropogenic preferences (Ross-Ibarra et al. 2007). In the case of plants, the obvious consequences of domestication are represented by visually recognizable traits that include changes in photoperiod response, reduced capacity of seed dispersion, reduced lateral branching, larger fruits or grains, reduced duration of seed dormancy, and increase in the size of female inflorescences (Pickersgill 2007; Larson et al. 2014). Notably, research efforts to understand the phenotypic changes that gave rise to maize have mainly focused on the direct consequences of domestication in aerial organs and sometimes roots (Chen et al. 2022; Lopez-Valdivia et al. 2022; Ren et al. 2022), with no emphasis on the role of adaptive environmental responses that, by acting on specific gene function, could have given rise to phenotypic traits that were identified and selected by humans.

It is generally accepted that domestication of maize (*Zea mays* ssp. *mays*) from teosinte *parviglumis* (*Zea mays* ssp. *parviglumis*) initiated in Mesoamerica around 9,000 years before present (Matsuoka et al. 2002). The initial evolutionary transition that resulted in the event of subspeciation most likely occurred close to the Balsas river drainage, in regions geographically converging with the Trans-Mexican Volcanic Belt (TMVB), at the intersection of the States of Mexico, Guerrero, and Michoacan (Matsuoka et al. 2002). Although the nature of the initial phenotypic changes that allowed the emergence of primitive maize remains unclear, the most obvious architectural and morphological differences between both subspecies affect their vegetative and reproductive organs. Teosinte *parviglumis* generates numerous lateral branches ending in male inflorescences, while maize often has a few short or no lateral branches, not ending in male reproductive organs (Wills et al. 2013). Additionally, the seeds of teosinte *parviglumis* are covered by a hard glume that protects the kernel externally, while in maize glumes are reduced, soft and do not cover the kernel (Doebley and Stec 1993). Large scale efforts involving physical mapping and comparative genomic analysis have resulted in the identification of numerous regions showing differential nucleotide variability between teosinte *parviglumis* and the B73 reference maize genome. In some cases, mutations in genes located within those genomic regions result in phenotypes that reveal their involvement in domestication traits. One of the major genes affected by maize domestication is *Teosinte branched1* (*Tb1*), a transcription factor of the TCP family that controls lateral branching, sexual determination, and glume formation (Doebley et al. 1997; Martín-Trillo and Cubas 2010). *Tb1* acts as a repressor of lateral branching (Studer et al. 2017), giving rise to maize plants usually composed of a single stem. Other genes involved in domestication include *Teosinte glume architecture1 (Tga1)*, *ramosa2 (ra2)* and *Zea floricaula leafy2 (zfl2),* among others (Bomblies and Doebley 2006; Sigmon and Vollbrecht 2010; Wang et al. 2015).

The importance of phenotypic plasticity is considered as likely relevant in the domestication history of crops (Ross-Ibarra et al. 2007; Lorant et al. 2020; Mueller et al., 2023); however, the role of environmental factors influencing their origins remains elusive and poorly investigated. A previous study showed that teosinte *parviglumis* grown under low temperatures and CO_2_ concentrations - reminiscent of those prevailing in the late Pleistocene and early Holocene - exhibit some phenotypes related to the domestication syndrome, including short branches, a single main stalk, and partially open seed fruitcases (Piperno et al. 2015; Piperno et al. 2019); however, the genetic basis of these changes have not been investigated. Although these results suggest that phenotypic plasticity could have influenced the domestication process, to this date there are no hints on how the adaptation to local environmental conditions could have affected specific gene function and guided human selection.

Recent genomic evidence indicates that admixture of maize with native teosinte *mexicana* populations (*Zea mays* ssp. *mexicana*) contributed to maize adaptation across a wide range of altitudes and latitudes in the American continent (Yang et al. 2023); however, the earliest maize macro-specimens found to date lack teosinte *mexicana* admixture and exhibit morphological attributes of fully domesticated cobs, indicating that *mexicana* is not a direct ancestor of maize and that its introgression occurred after initial divergence from teosinte *parviglumis* (Vallebueno-Estrada et al. 2023; Acosta-Bayona et al. 2024). Consequently, the nature of the genetic factors that guided the emergence of maize remain largely unknown, and the initial phenotypic changes that caused the original divergence of these two subspecies have not been identified. It is also unknown if domestication initiated by direct action of human selection on teosinte natural standing variation, or if the response to specific environmental factors generated phenotypic changes that were identified and selected by humans to act on native teosinte *parviglumis* populations.

Maize can adapt to abiotic factors such as heavy metal (HM) stress through numerous mechanisms that prevail during development (AbdElgawad et al. 2020; Gallo-Franco et al. 2020; Malenica et al. 2021; Gao et al. 2022). These include metal immobilization by mycorrhizal association, metal sequestration, or complexation by exuding organic compounds from roots, metal intracellular sequestration and compartmentalization in vacuoles. HMs include essential elements that are necessary for maize growth through enzymatic function, such as iron (Fe), manganese (Mn), zinc (Zn), magnesium (Mg), molybdenum (Mo), and copper (Cu); and non-essential elements like Cadmium (Cd), chromium (Cr), lead (Pb), aluminum (Al), and selenium (Se). In general, HM stress leads to visibly reduced plant growth due to the reduced cell elongation and cell wall elasticity. At the molecular level, metal toxicity can disturb the cellular redox balance and lead directly or indirectly to oxidative stress and the formation of reactive oxygen species (ROS), shifting the redox balance to the pro-oxidative side until the antioxidative defense system allows a new balanced redox status (Smeets et al. 2009; Hall 2002).

The comparison of the *Palomero toluqueño* and B73 genomes resulted in the identification of a large collection of identical sequence regions (IdSRs) with low nucleotide variability and transcriptional evidence of expression (Vielle-Calzada et al. 2009). Putative domestication loci included *ZmHMA1* and *ZmHMA7*, two genes encoding members of the P1_b_-ATPases protein superfamily, also known as Heavy Metal ATPases (P-type ATPases) characterized by the generation of a phosphorylated intermediate during the transport reaction cycle. Members of this superfamily contain 8 to 12 transmembrane domains as well as conserved ATP binding, phosphorylation and dephosphorylation sites (Solioz and Vulpe 1996; Axelsen and Palmgren 2001; Voskoboinik et al. 2003; Williams and Mills 2005). The superfamily can be divided in subfamilies that include H±ATPases (type 3_A_) in plants and fungi, Na+/K ATPases (type 2_C/D_) in animals, Ca2±ATPases (type 2_A/B_), and heavy metal (HM) transporting ATPases (type 1_B_) capable of transporting a large variety of HM cations across cell membranes. *ZmHMA1* and *ZmHMA7* are type 1_B_ ATPases that include a conserved Cys-Pro-Cys/His/Ser motif within a transmembrane domain necessary for the translocation of HMs cations through the membrane channel. A third *Palomero toluqueño*/B73 IdSR included *Skewed Root Growth Similar5* (*ZmSKUs5*), a gene encoding a Skewed5 (SKU5)-similar (SKS) protein classified as a multi-copper oxidase containing cuperdoxin domains (Sedbrook et al. 2002). All three genes map to chromosome five (chr.5), with *ZmHMA1* and *ZmSKUs5* within a genomic region that fractionates into multiple QTLs under selection during the *parviglumis* to maize transition (Doebley and Stec 1993; Lemmon and Doebley, 2014). Contrary to major domestication loci showing a single highly pleiotropic gene responsible for important domestication traits, in this chr.5 genomic region the smallest 1.5-LOD support interval contained more than 50 genes, suggesting that phenotypic effects are due to multiple linked QTLs (Lemmon and Doebley, 2014). While the functional characterization of all three maize genes remains elusive, the Arabidopsis and barley *ZmHMA1* homologs act during heavy metal transport in the chloroplast (Boutigny et al. 2014), demonstrating their functional involvement in homeostasis.

To investigate the possibility that HM response had an effect on the origin and domestication of maize, we compared the phenotypic changes that teosinte *parviglumis* and maize undergo when grown in the presence of sublethal concentrations of Cu and Cd. We show for the first time that teosinte *parviglumis* responds to HM stress by acquiring an aerial architecture reminiscent of extant maize and unusual proliferation of female inflorescences. *Tb1* expression is upregulated in teosinte *parviglumis* meristems when grown under HM stress. Analysis of nucleotide variability across 11.46 Mb of chr.5 indicates that many genes within the *ZmHMA1 to ZmSKUs5* genomic region show reduced nucleotide variability in maize as compared to teosinte *parviglumis*. Although the reduction is not specific to these genes, large-scale transcriptional analyses suggests that contrary to genes involved in other types of abiotic stress, a large number of loci that contain HM response genes were affected by positive selection during the teosinte *parviglumis* to maize transition. Both *ZmHMA1* and *ZmHMA7* are active throughout maize and teosinte *parviglumis* development, and their expression responds to HM stress. In addition, mutations in *ZmHMA1* show domestication-related defects such as the absence of seminal roots and increased height, both manifested under exposure to HMs. Paleoenvironmental studies reveal a temporal and geographical convergence that relates the history of volcanic eruptions in the TMVB with the emergence of maize at times of climatic instability. Overall, this evidence indicates that the evolutionary transition that gave rise to extant maize was influenced by heavy metal stress through the activity of P1b-ATPases, presumably responding to edaphological consequences of volcanic activity.

## Results

### Teosinte parviglumis responds to heavy metal stress by acquiring an architecture reminiscent of extant maize

To compare HM response in teosinte *parviglumis* and maize, we exposed wild-type W22 maize inbred and CIMMYTMA 29791 teosinte *parviglumis* individuals to sublethal concentrations of Cu (400 mg/kg of CuSO_4_) and Cd (16mg/kg of CdCl_2_). CuSO_4_ and CdCl_2_ were homogenized in soil before transplant and grown under controlled greenhouse conditions. HM concentrations were based on preliminary dose responses that allowed maize plants to reach reproductive stages and complete their life cycle (AbdElgawad et al. 2020). Whereas higher doses impair flowering or are lethal, lower Cu/Cd concentrations do not consistently show conventional phenotypic responses such as reduced plant growth (AbdElgawad et al. 2020; Atta et al., 2023). As previously reported (Atta et al. 2023), maize individuals grown in the presence of HMs showed a significant reduction in aerial traits such as stem diameter, size of leaves, size and number of roots, and size of female inflorescences (Figure 1; Tables 1 and S1). On average, stem diameter showed a significant reduction of about 17% (P<0.05) as compared to plants grown in absence of HM stress (Figure 1f to 1j). Similarly, the length and width of developing leaves showed significant reductions of 12%, and mature leaves of 6% (P<0.05). Reproductive traits such as the number of female inflorescences (NFI) and their average weight (WFI) showed reductions, as previously reported (NFI = 1.20 ± 0.42 in absence of HMs and NFI = 0.70 ± 0.67 in presence of HMs; WFI = 40.06 ± 14.39 in absence of HMs and WFI = 37.75 ± 18.83 in presence of HMs; Table 1). However, the most significant effects occurred in the root system, with an increase in the number of brace roots under HMs (NBR = 12.90 ± 4.33 in absence of HMs; NBR = 16.30 ± 4.99 in presence of HMs; Table 2), and a significant decrease in their length (LBR = 16.15 ± 9.75 cm in absence of HMs; LBR = 4.00 ± 2.04 cm in presence of HMs; P<0.001; Table 1). The number and length of crown roots were reduced on average by 33% and 18%, respectively (NCR = 11.30 ± 2.11 in absence of HMs; NCR = 7.60 ± 0.97 in presence of HMs; LCR = 58.90 ± 21.87 cm in absence of HMs; LCR = 48.85 ± 11.80 cm in presence of HMs; Table 1). Nodal roots also showed a significant reduction in their number and length (P<0.01 and P<0.001, respectively; NNR = 10.50 ± 2.27 in absence of HMs; NNR = 6.70 ± 2.79 in presence of HMs; LNR = 87.20 ± 16.80 cm in absence of HMs; LNR = 55.30 ± 21.10 cm in presence of HMs; Table 1). By contrast, the number (NSR) and length (LSR) of seminal roots was significantly increased in the presence of HMs (P<0.001 and P<0.01, respectively: NSR = 4.10 ± 1.45 in absence of HMs; NSR = 6.60 ± 1.35 in presence of HMs; LSR = 27.70 ± 13.38 cm in absence of HMs; LSR = 43.50 ± 14.01 cm in presence of HMs) when measured 50 days after transplant (Table 1).

**Figure 1.**
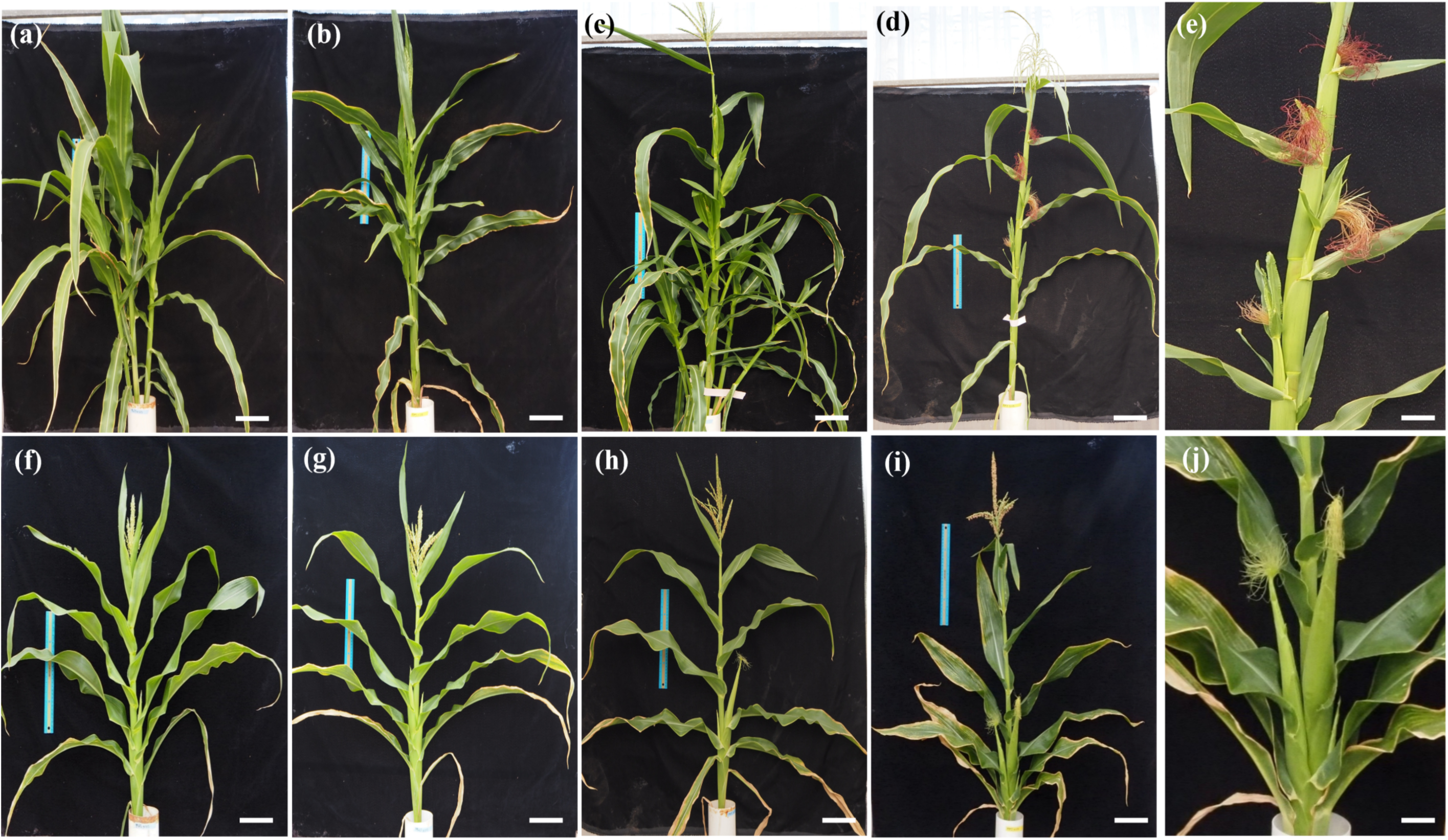
Phenotypic response of teosinte *parviglumis* and wild-type W22 maize grown under heavy metal stress. (**a**) to (**e**), teosinte *parviglumis*; (**f**) to (**j**), W22 maize. (**a**) Teosinte *parviglumis* in absence of heavy metal stress at early flowering stage. (**b**) Teosinte *parviglumis* grown under heavy metal stress at early flowering stage. (**c**) Teosinte *parviglumis* in absence of heavy metal stress at late flowering stage. (**d**) Teosinte *parviglumis* grown under heavy metal stress at late flowering stage. (**e**) Teosinte *parviglumis* showing absence of lateral branching, and proliferation of female inflorescences arising from independent axillary meristems under heavy metal stress (detail of d). (**f**) W22 maize in absence of heavy metal stress at early flowering stage. (**g**) W22 maize grown under heavy metal stress at early flowering stage. (**h**) W22 maize in absence of heavy metal stress at late flowering stage. (**i**) W22 maize grown under heavy metal stress at late flowering stage. (**j**) W22 maize female inflorescences at late flowering stage (detail of i). Scale bars: 10 cm.

**Table 1.**
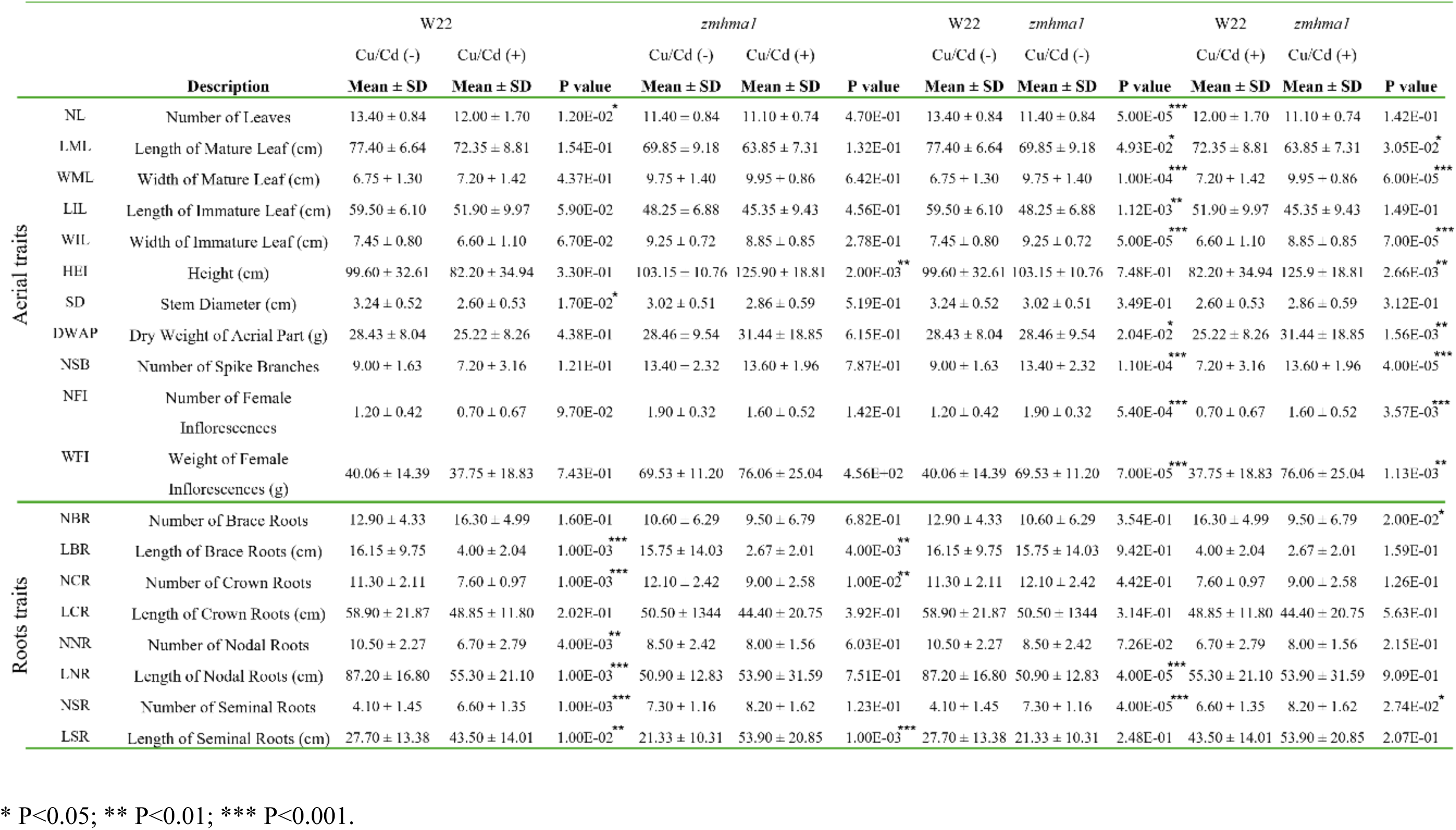
Estimation of phenotypic traits in wild-type and z*mhma1* maize grown in absence and presence of heavy metal stress.

**Table 2.**
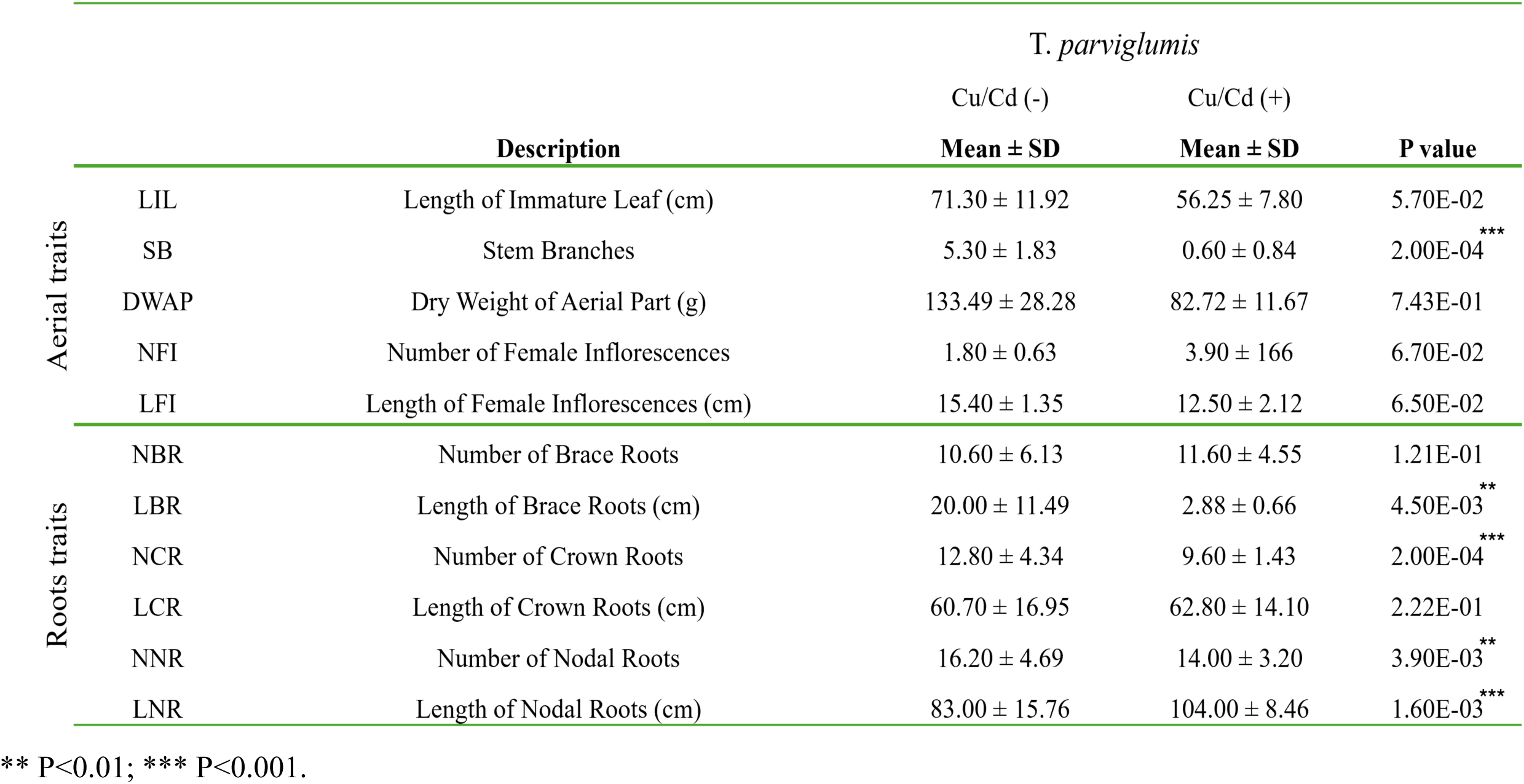
Estimation of phenotypic traits in teosinte *parviglumis* grown in absence and presence of heavy metal stress.

Interestingly, teosinte *parviglumis* exhibited a maize-like phenotype when grown in the presence of HMs, with complete absence of tillering in most plants (Figure 1; Tables 2, S1 to S3). This phenotype had never been reported for teosinte *parviglumis* plants grown under HM stress or any other type of edaphological conditions. From a total of 30 plants tested, 28 showed total absence of tillering, and two initiated secondary stems during vegetative stages but arrested their growth early during development, resulting in the presence of a single stem at reproductive stages (Figure 1a to 1d; and Table 2). Additionally, all 30 plants tested showed complete absence or residual lateral branching with unusual proliferation of female inflorescences arising from independent axillary meristems in primary stems (Figure 1e). This additional phenotype had not been reported for teosinte *parviglumis* under any type of environmental conditions. The increase in the number of female inflorescences clustered within a single stem (NFI = 1.80 ± 0.63 in absence of HMs; NFI = 3.90 ± 1.66 in presence of HMs; Figure 1e and Table 2) was associated with shorter flowering time as compared with normal growing conditions in absence of HMs (Figure 1d). In addition, teosinte *parviglumis* grown in the presence of HMs showed reduced leaf length (LIL = 71.30 ± 11.32 cm in absence of HMs; 56.25 ± 7.80 cm in presence of HMs; Table 2). The absence of lateral branches resulted in lower values of aerial dry weight (DWAP = 133.49 ± 28.28 gr in the absence of HMs; DWAP = 82.72 ± 11.67 gr in presence of HMs; Table 2). Teosinte *parviglumis* grown under HMs showed a significant reduction in the length of brace roots (P<0.01; LBR = 20.00 ± 11.49 cm in absence of HMs; LBR = 2.88 ± 0.66 cm in presence of HMs; Table 2), but no significant changes in their number. The number crown and nodal roots was significantly decreased by HM stress (P<0.001 and P<0.01 respectively; NCR=12.88 ± 4.34 in absence of HMs; 9.6 ± 1.43 in presence of HMs; NNR=16.2 ± 4.69 in absence of HMs; NNR= 14 ± 3.2 in presence of HMs; Table 2). By contrast, the length of nodal roots was significantly increased by HM stress (P<0.001; LNR=83 ± 15.76 in absence of HMs; 104 ± 8.46 in presence of HMs). The presence of HMs did not modify the absence of seminal roots intrinsic to teosinte *parviglumis* accessions (Perkins and Lynch 2021). Taken together, these results suggest that the response of maize and teosinte *parviglumis* to HM stress can give rise to phenotypes previously shown to distinguish both subspecies at the light of the domestication syndrome.

### Genomic regions encompassing ZmHMA1, ZmHMA7 and ZmSKUs5 show reduced nucleotide variability in extant maize as compared to teosinte parviglumis

*ZmHMA1* and *ZmSKU5* mapped 52.1 and 41.8 cM, respectively, in a genetic map of chr.5 calculated to be 86.64 cM, with an average megabase pair to centimorgan ratio of 1.873 Mbp/cM (Lemmon and Doebley, 2014; Figure S1). The corresponding mapping population used isogenic recombinant inbred lines (NIRILs) with informative cross-over events concentrated in the region of interest and replicated phenotypic assessments in block experiments. Multiple QTLs were detected in a region spanning across 82 cM, none with single large effect genes such as those found in other chromosomes (Wang et al., 2005; Studer et al. 2011; Wills et al., 2013), suggesting that several associated domestication traits are due to multiple linked QTLs of small effect (Lemmon and Doebley, 2014). The location of *ZmSKU5* overlaps with a single wide 1.5-LOD support interval that spreads across 50.6 cM (number of kernels per rank; Figure S1). By contrast, *ZmHMA1* overlaps with a contiguous region containing no less than five QTL 1.5- LOD support intervals (ear diameter; number of kernels per rank; percentage of staminate spikelets; tassel branch number; and tillering index; Figure S1).

To determine if the regulatory and coding regions of *ZmHMA1, ZmHMA7,* and *ZmSKUs5* were affected by human selection, we calculated their level of nucleotide variability based on single nucleotide polymorphisms (SNPs) for all extant maize and teosinte *parviglumis* accessions included in the diversity panel HapMap3, using the corresponding sequences of *Tripsacum dactyloides* as an outgroup. The levels of the nucleotide variability index - π(m) and π(t) for maize and teosinte *parviglumis*, respectively - were determined in bins of 100 nt using VCFtools, including alignments encompassing a genomic region that included 15 kb upstream of the START codon, the corresponding coding region for each gene, and 15 kb downstream of the STOP codon (Figure 2 and Table S4). In the case of *ZmHMA1*, the upstream region values were π(m)= 0.00071 and π(t)= 0.00870, and for the downstream region, π(m)=0.00097 and π(t)=0.00248. By contrast, the *ZmHMA1* coding sequence showed low values in both subspecies: π(m)= 0.0050 and π(t)= 0.00106. These results indicate that human selection in the *ZmHMA1* locus preferentially affected its upstream and downstream regulatory regions (Figure 2a and Table S4). *ZmHMA7* showed contrasting values of π(m) and π(t) across the entire coding sequence and its regulatory regions (Figure 2b). For the upstream region, values were π(m)= 0.0009 and π(t)=0.00411; for the *ZmHMA7* coding region, π(m)= 0.00015 and π(t)= 0.00148; and for the downstream region, π(m)=0.00066 and π(t)=0.00454. *ZmSKUs5* also showed contrasting nucleotide variability in the entire coding and flanking regions (Figure 2c and Table S4). Values were π(m)= 0.0011 and π(t)= 0.00735 for the upstream region, π(m)= 0.00044 and π(t)= 0.00754 for the coding sequence, and π(m)=0.00203 and π(t)=0.00626 for the downstream region (Table S4). For all three genes, similar π values in the coding region of maize landraces and improved lines indicates that the loss of nucleotide variability occurred during the teosinte *parviglumis* to maize transition, and not during subsequent contemporary breeding (Table S4). By contrast, the *ZmGLB*1 neutral control locus shows equivalent levels of nucleotide variability in maize and teosinte *parviglumis* (Eyre-Walker et al. 1998).

**Figure 2.**
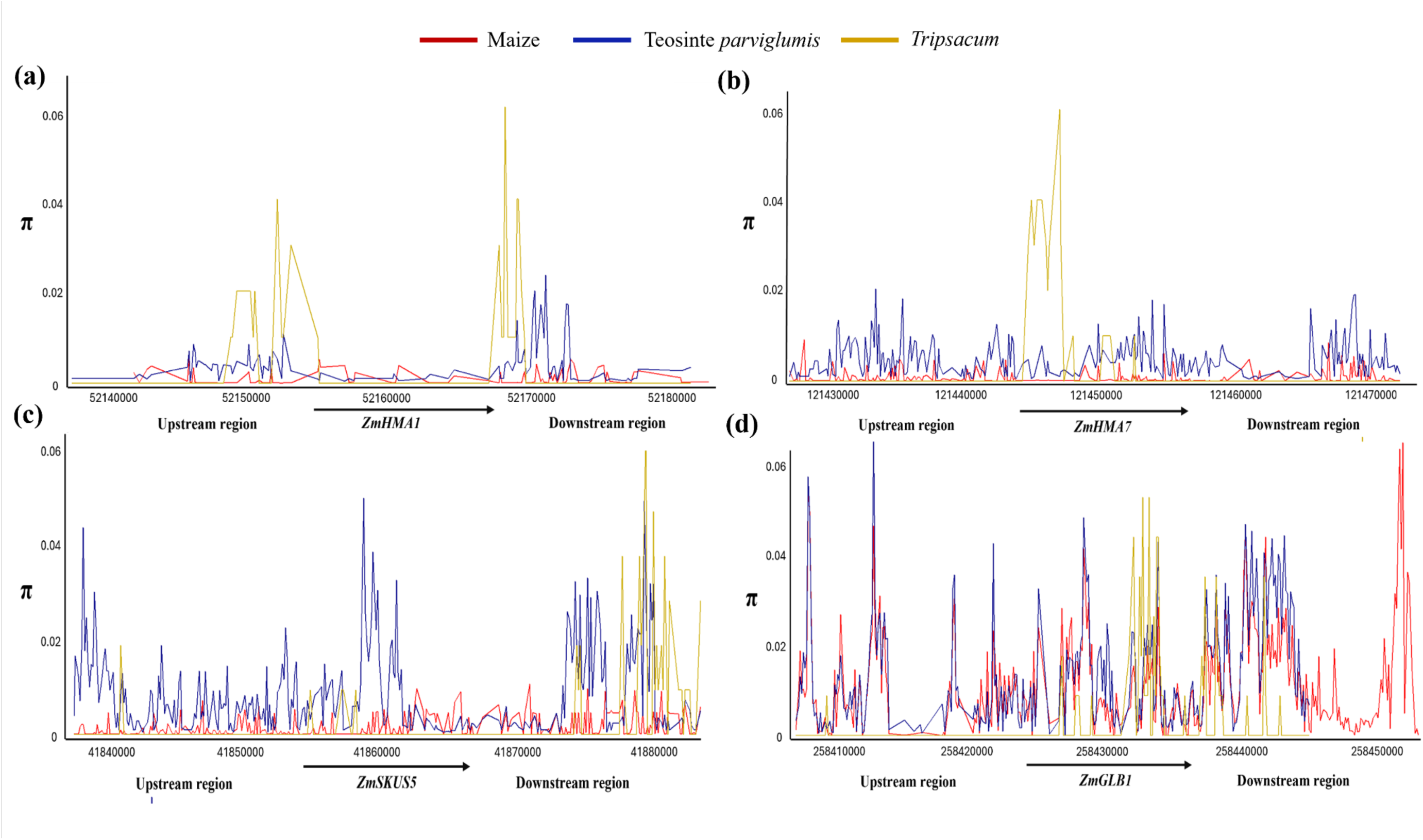
Genetic diversity of loci encompassing *ZmHMA1, ZmHMA7*, and *ZmSKUs5.* The nucleotide variability index (π) was calculated for all HapMap3 accessions of teosinte *parviglumis* (blue line), maize (red line), and *Tripsacum dactyloides* (orange line), taking in consideration the coding sequence (arrow) and an upstream and downstream region of 15 Kb encompassing each gene. (**a**) Nucleotide variability in the *ZmHMA1* locus. (**b**) Nucleotide variability in the ZmHMA7 locus. (**c**) Nucleotide variability in the *ZmSKUs5* locus. (**d**) Nucleotide variability in the *ZmGLB1* locus.

We also compared the level of genetic variability in *ZmHMA1*, *ZmSKUs5*, and *ZmHMA7* (and their neighboring regions) to all chr.5 genes categorized as likely affected by domestication (Hufford et al., 2012), and to the full collection of neutral genes distributed across the whole chromosome. The analysis was conducted using 100 bp bins and sliding windows of 50 bp; for a total of 385,829 bins in the case of chr.5 domestication genes, and 2,373,010 bins for neutral genes. As shown in Figure S2, the levels of nucleotide variability in the three candidate genes and their six neighboring loci are significantly reduced in maize landraces as compared to teosinte *parviglumis* inbred lines. When compared to the full neutral gene set of chr.5, this reduction is more severe than the reduction affecting 304 previously identified domestication genes, confirming the exceptional selective pressure that was imposed in those three loci during the evolutionary transition from teosinte *parviglumis* to extant maize.

### Large-scale genomic and transcriptomic comparisons indicate that many HM response genes were positively selected across the maize genome

To expand the results well beyond the analysis of the three genes previously described, we performed a detailed analysis of genetic diversity across the 11.47 Mb genomic region comprised between Z*mSKUs5* and *ZmHMA1*. This additional analysis reveals general tendencies in the quantity and nature of loci that were affected by positive selection during the teosinte *parviglumis* to maize transition in region identified via LOD score on chr.5. We compared nucleotide variability by using 100 bp bins covering loci composed of two 30 Kb segments up and downstream of coding sequences, respectively, and the coding sequence itself, for 173 genes present within the genomic region comprised between *ZmSKUs5* and *ZmHMA* (Figure S1 and Supplementary File 6). Two types of statistical tests (ANOVA and Wilcoxon) were applied to nucleotide variability comparisons using the entirety of each locus. The Benjamini-Hochber procedure allowed an estimation of the false discovery rate (FDR<0.05) to avoid type I errors (false positives). Although some individual loci appear as differently classified depending on the statistical test applied (22 out of 173 loci), the general differences in nucleotide variability are consistently maintained within the subregions described below. We found that 166 out of 173 loci show signatures of positive selection and are roughly organized in five independent subregions of variable length. The first six loci are consecutively ordered in a 402 Kb subregion that includes *ZmSKUs5.* A second group of 13 consecutive loci expands over a 1.44 Mb subregion that contains *NRAMP ALUMINUM TRANSPORTER1*, also involved in HM response through uptake of divalent ions. A third group of 17 consecutive loci expands over 1.28 Mb; eleven contain genes encoding for uncharacterized proteins. The fourth group is composed of 57 consecutive loci expanding over 3.22 Mb and contains genes encoding for *DEFECTIVE KERNEL55, AUXIN RESPONSE FACTOR16*, and peroxydases involved in responses to oxydative stress. The fifth group contains 12 consecutive loci expanding over 713 Kb and contains *ZmHMA1.* An additional segment of approximately 1.17 Mb and containing 25 consecutive loci that were positively selected expands away from the *ZmSKUs5-ZmHMA1* segment; it also contains several genes encoding for peroxydases. Although multiple loci include genes that could be involved in abiotic stress and oxidative responses, these results suggest that multiple factors other than HM stress could have played a role in the evolutionary mechanisms that affected the genetic diversity of chr.5 during the teosinte *parviglumis* to maize transition.

To further analyze the possibility that HM response could have played a role in maize emergence and subsequent domestication, we analyzed large scale transcriptomic data corresponding to independent experiments aiming at understanding the response of maize roots to HM stress. Six available transcriptomes were selected for in-depth analysis because they presented a fold change strictly higher than 1, and their results were supported by false discovery rates (FDR<0.05). These six transcriptomes (Table S5) included HM response datasets corresponding to growth conditions that not only incorporated Cu, but also lead (Pb) and chromium (Cr) that were not included in the substrate of our experiments. Transcriptional profiles were obtained from roots of plants at different stages: maize seedlings (Shen et al., 2012; Gao et al., 2015; Zhang et al., 2024a), three week old plantlets (Yang et al., 2023), and plants at V2 stage (Zhang et al., 2024b; Fengxia et al., 2025). A total of 120 genes shared by all six transcriptomes were found to be differentially expressed under HM stress conditions (66 upegulated and 54 downregulated; Figure S3), including *ZmSKUs5*, *ZmHMA1* and *ZmHMA7*; 52 of them (43.3%) are located in maize loci showing less than 70% of the nucleotide variability found in teosinte *parviglumis*, suggesting that they were affected by positive selection (Yamasaki et al., 2005; Supplementary File 7). Of 18 mapping in chr.5, twelve are within the 82 cM that fractionates into multiple QTLs under selection during the *parviglumis* to maize transition. Interestingly, five additional loci containing HM response genes completely lack SNPs within their total length in both *parviglumis* and maize, and 19 additional loci lack SNPs in at least one 30 Kb segment or their coding region (Supplementary File 7), suggesting the frequent presence of ultraconserved genomic regions in many loci containing HM response genes. When this same analysis was conducted in a set of loci comprising 63 genes previously identified as differentially expressed in response to abiotic stress not directly related to HM responses (hypoxia; nutritional deficiency; soil alkalinity; drought; soil salinity), 18 loci (28.6%) showed less than 70% of the nucleotide variability found in teosinte *parviglumis*. Only one of them maps in chr.5 and none contained segments or coding regions lacking SNPs in *parviglumis* or maize. These results suggest that in contrast to other types of abiotic stress response genes, loci comprising a large set of genes that unambiguously respond to HM stress caused by chemical elements of diverse nature were affected by positive selection during the *parviglumis* to maize transition, irrespectively of their position in the genome.

### ZmHMA1 and ZmHMA7 are active throughout development and their expression is influenced by the presence of heavy metals

The gene structure of *ZmHMA1, ZmHMA7,* and *ZmSKUs5* is shown in Figure 3. The coding sequence of *ZmHMA1* and *ZmHMA7* is approximately 21.6 Kb and 4.5 Kb, respectively, while *ZmSKUs5* is approximately 4.2 Kb in length. Within the short arm of chromosome 5, *ZmHMA1* and *ZmSKUs5* are separated by 9.16 Mb, with *ZmHMA1* approximately 15 Mb away from the maximum likelihood prediction of a QTL involved in maize domestication (Doebley and Stec 1993; Vielle-Calzada et al. 2009). *ZmHMA7* maps to the centromeric region of chromosome 5. A detailed PlantCARE-based analysis identified several conserved binding sites and regulatory elements in the promoter region of these three genes (Lescot et al. 2002). *ZmHMA1* and *ZmHMA7* share binding sites to basic Helix-Loop-Helix (bHLH) transcription factors, as well as abscisic acid response (ABRE), ethylene responsive (ERF), and G-box (CACGTG) elements recognized by leucine zipper family (bZIP) and bHLH transcription factors (Figure 3a). The regulatory region of both *ZmHMA1* and *ZmSKUs5* contain a binding site for MIKC-type MADS box transcription factors (Figure 3a), whereas *ZmHMA7* and *ZmSKUs5* share a jasmonic acid-response TGACG-motif (Figure 3a), as well as a Myb-binding (MBS) and GATA binding site. In addition, *ZmHMA1* contains a binding site for TCP proteins such as Tb1, increasing the number of families of transcription factors that could be involved in shared or individual regulation of these three genes (Figure 3a).

**Figure 3.**
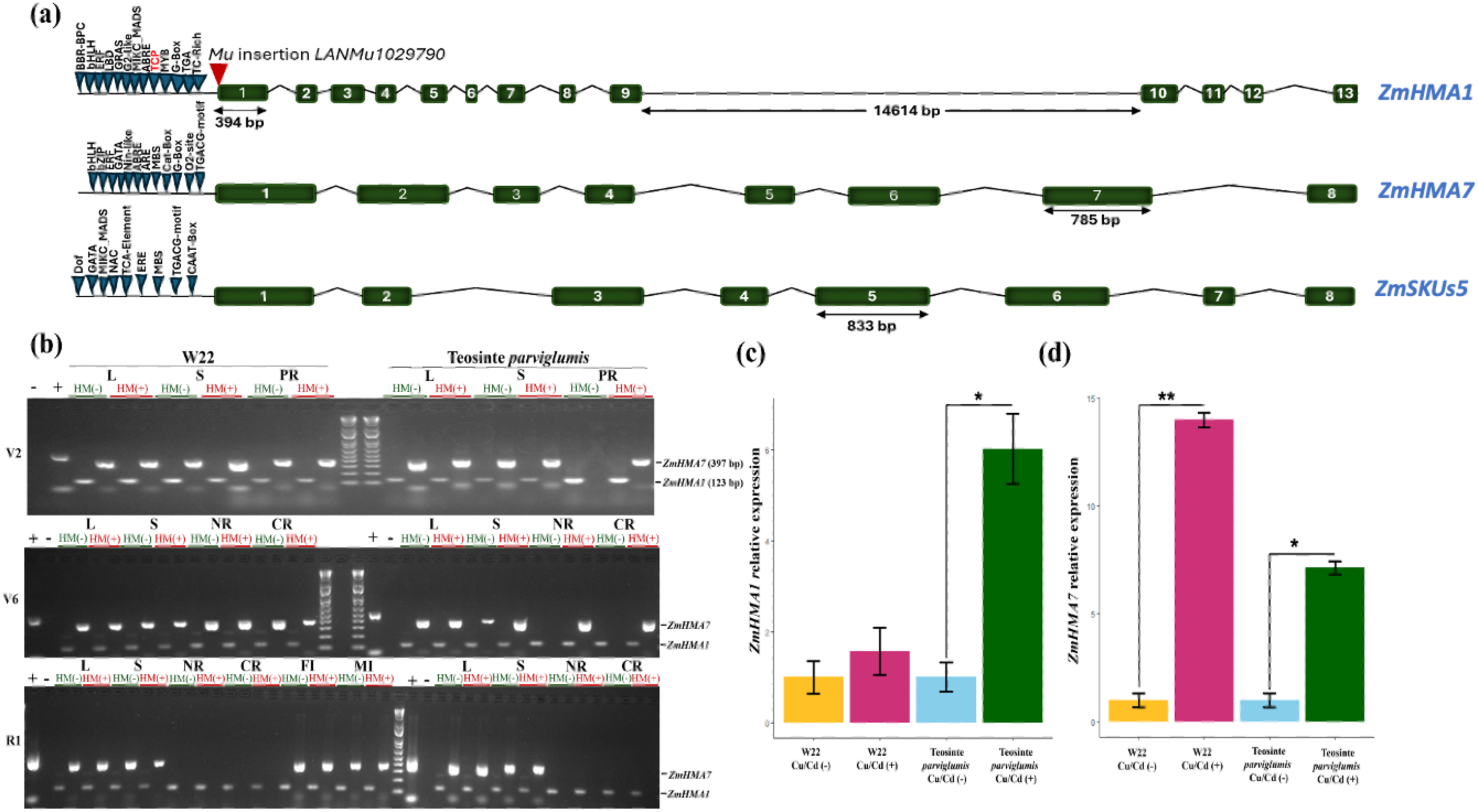
Structure and gene expression of *ZmHMA1* and *ZmHMA7* in teosinte *parviglumis* and W22 maize. (**a**) Structural analysis of *ZmHMA1*, *ZmHMA7*, and *ZMSKUs5*. Blue triangles indicate the location of motifs predicted in the regulatory region of each gene. The insertion site of *Mu* line *LANMu1029790* is indicated by a red triangle. (**b**) RT-PCR analysis of *ZmHMA1* and *ZMHMA7* expression throughout development of teosinte *parviglumis* and W22 maize in presence or absence of heavy metal stress; L=leaf; S=stem; PR=primary root; NR=nodal root; CR=crown root. V2: 15 days after transplant; V6: 30 days after transplant; R1: 50 days after transplant. Controls: (-) absence of DNA; (+) genomic DNA. (**c**) qPCR analysis of *ZmHMA1* expression in 1-month old primary roots, in absence or presence of heavy metal stress. (**d**) qPCR analysis of *ZmHMA7* expression in 1-month old primary roots, in absence or presence of heavy metal stress. Histograms in (**c**) and (**d**) show the relative fold-change (normalized to actin, based on technical and biological triplicates); standard deviations are shown by vertical lines and Student T-test P-values represented by asterisks. * P<0.01; ** P<0.001.

To elucidate the expression patterns of *ZmHMA1* and *ZmHMA7*, we conducted reverse transcriptase PCR (RT-PCR) in leaves, stems, roots, and flowers at two vegetative (V2 and V6) and one reproductive (R1) developmental stage, in teosinte *parviglumis* and wild-type W22 maize individuals (Figure 3b). In the case of maize, *ZmHMA1* was expressed in leaves, stems, roots and flowers throughout development in either absence or presence of HMs. *ZmHMA7* was also expressed in aerial organs throughout development in both absence and presence of HMs but was not expressed in roots at reproductive stage R1. In teosinte *parviglumis, ZmHMA1* was also expressed throughout development in either absence or presence of HMs. By contrast, ZmHMA*7* was constitutively expressed in aerial organs, but only expressed in roots under HM stress during V2 and V6 stages. To further determine if the level of expression of these two genes is influenced by the presence of HMs, we conducted quantitative real-time PCR (qPCR) using the primary root of teosinte *parviglumis* and maize plantlets at four weeks after germination (Figures 3c and 3d). In wild-type W22 maize, the expression of *ZmHMA1* was not significantly different in the presence and absence of HMs. By contrast, in teosinte *parviglumis* the expression of *ZMHMA1* was significantly increased in presence of HMs (Figure 3c). In the case of *ZmHMA7*, expression was significantly increased in both maize and teosinte *parviglumis* when plants were grown in the presence of HMs (Figure 3d). Overall, these results indicate that the activity of both *ZmHMA1* and *ZmHMA7* is influenced by HM stress in teosinte *parviglumis* and maize.

### ZmHMA1 null mutant individuals show phenotypic defects related to the domestication syndrome

Transposon *Mu* insertional lines have been widely used to elucidate gene function in maize. To elucidate the function of *ZmHMA1*, we phenotypically analyzed *LANMu1029790*, a 4-generation backcross of line *mu1029790* into the W22 background selected for single copy insertions in *ZmHMA1* (McCarty et al. 2005). In *LANMu1029790,* a single *Mu* insertion is in the first exon of *ZmHMA1*, 65 bp downstream of the START codon (Figure 3a). RT-PCR analysis showed that expression of Z*mHMA1* is completely absent in homozygous individuals for the *LANMu1029790* insertion (Figure S4), confirming that the corresponding mutant is a null allele. Plants harboring a homozygous *LANMu1029790* insertion are subsequently designated as *zmhma1-m1::MuDR* (abbreviated *zmhma1-m1*).

To determine the function of *ZmHMA1* in maize, we compared the phenotype of wild-type W22 and homozygous *zmhma1-m1* individuals at the reproductive R1 stage (Table 1). Under normal growing conditions, homozygous *zmhma1-m1* individuals generated significantly less leaves than wild-type (P<0.001; NL = 13.4 ± 0.84 in wild-type, and NL = 11.4 ± 0.84 in *zmhma1-m1*). Whereas the length of mutant leaves was smaller than wild-type (LML = 77.4 ± 6.64 cm in wild-type, and LML = 69.85 ± 9.18 cm in *zmhma1-m1*), their width was significantly larger (P<0.001; WML = 6.75 ± 1.3 cm in wild-type, and WML = 9.75 ± 1.4 cm in *zmhma1-m1*), resulting in an increase of the total dry weight of aerial organs in mutant plants (DWAP = 18.43 ± 8.04 grams in wild-type, and DWAP = 28.46 ± 9.54 grams in *zmhma1-m1*). Homozygous *zmhma1*-*m1* individuals also produced significantly more female inflorescences per plant (P<0.001; NFI = 1.2 ± 0.42 in wild-type, and NFI = 1.9 ± 0.32 in *zmhma1*) a difference that reflected in the total dry weight of female inflorescences per plant (P<0.001; WFI = 40.06 ± 14.39 gr in wild-type and WFI = 69.5 ± 11.2 gr in *zmhma1*); and significantly more spikes per male inflorescence (P<0.001; NSB = 9 ± 1.63 in wild-type, and NSB = 13.4 ± 2.32 in *zmhma1-m1*). Interestingly, the general length of roots was significantly smaller in homozygous *zmhma1-m1* individuals, in particular nodal roots (P<0.001; LNR = 87.2 ± 16.8 cm in wild-type, and LNR = 50.9 ± 12.83 cm in *zmhma1-m1*). By contrast, mutant plants showed a significant increase in the number of seminal roots per plant (P<0.001; NSR = 4.1 ± 1.45 in wild-type, and NSR = 7.3 ± 1.16 in *zmhma1-m1*). When wild-type and mutant individuals were grown under HM stress, differences in the size of leaves, number of female inflorescences per plant, and number of spikes per male inflorescence were maintained. In addition, mutant individuals exposed to HM stress were significantly taller than wild-type (P<0.01; HEI = 82.2 ± 34.94 cm in wild-type, and HEI = 125.9 ± 18.81 cm in *zmhma1-m1*). In addition, total root and seminal root length were equivalent in wild-type and mutant individuals, mainly because the length of wild-type roots was significantly reduced under HM stress; however, the number of seminal roots was again significantly higher in mutant individuals (P<0.05; NSR = 6.6 ± 1.35 in wild-type, and NSR = 8.2 ± 1.62 in *zmhma1-m1*). These results indicate that in maize aerial organs, *ZmHMA1* is involved in promoting leaf growth, restricting the number of female inflorescences per plant, and restricting the number of spikes in male inflorescences. In underground organs, *ZmHMA1* is involved in promoting root growth and restricting the number of seminal roots per individual. Under HM stress, *ZmHMA1* shows an additional role in restricting plant height, a function that was not identified under normal growing conditions. Overall, these results suggest that *ZmHMA1* is involved in the control of some of the aerial and underground phenotypic traits that distinguish maize from teosinte *parviglumis,* especially under HM stress conditions.

### Under heavy metal stress, Tb1 is overexpressed in the apical meristem of teosinte parviglumis

The absence of secondary ramifications in teosinte *parviglumis* grown under HM stress is reminiscent of the aerial architecture of wild-type maize. Previous QTL and molecular analysis suggested that *Tb1* contributed to the architectural difference between maize and teosinte, as *tb1* maize mutants develop secondary stem ramifications ending in male inflorescences, resembling teosinte in their overall architecture (Doebley et al. 1997). Interestingly, the regulatory region of *ZmHMA1* contains a specific binding site for members of the Teosinte-branched 1/Cycloidea/Proliferating (TCP) family of transcription factors that includes Tb1 (Figure 3a). To investigate the possibility that the phenotypic response of teosinte *parviglumis* under HM stress could be regulated by *Tb1*, we conducted quantitative real-time PCR (qPCR) to determine the expression of *ZmHMA1* and *Tb1* in the shoot apical meristem (SAM) of one month old W22 maize and teosinte *parviglumis* plantlets (Figure 4a and 4b). The expression *ZmHMA1* was downregulated in W22 wild-type maize meristems in the presence of HMs. By contrast, HM stress resulted in a significant increase of *ZmHMA1* expression in teosinte *parviglumis*, confirming that in the maize ancestor the level of *ZmHMA1* activity is influenced by the presence of HMs in the soil. In addition, the expression of *Tb1* was significantly upregulated in teosinte *parviglumis* meristems in the presence of HMs. These results do not allow to directly link the regulation of *ZmHMA1* expression to the function of *Tb1*; however, they open an opportunity to further investigate the possibility that under HM stress, the formation of secondary ramifications in teosinte *parviglumis* could be repressed by transcription factors of the TCP family, including Tb1.

**Figure 4.**
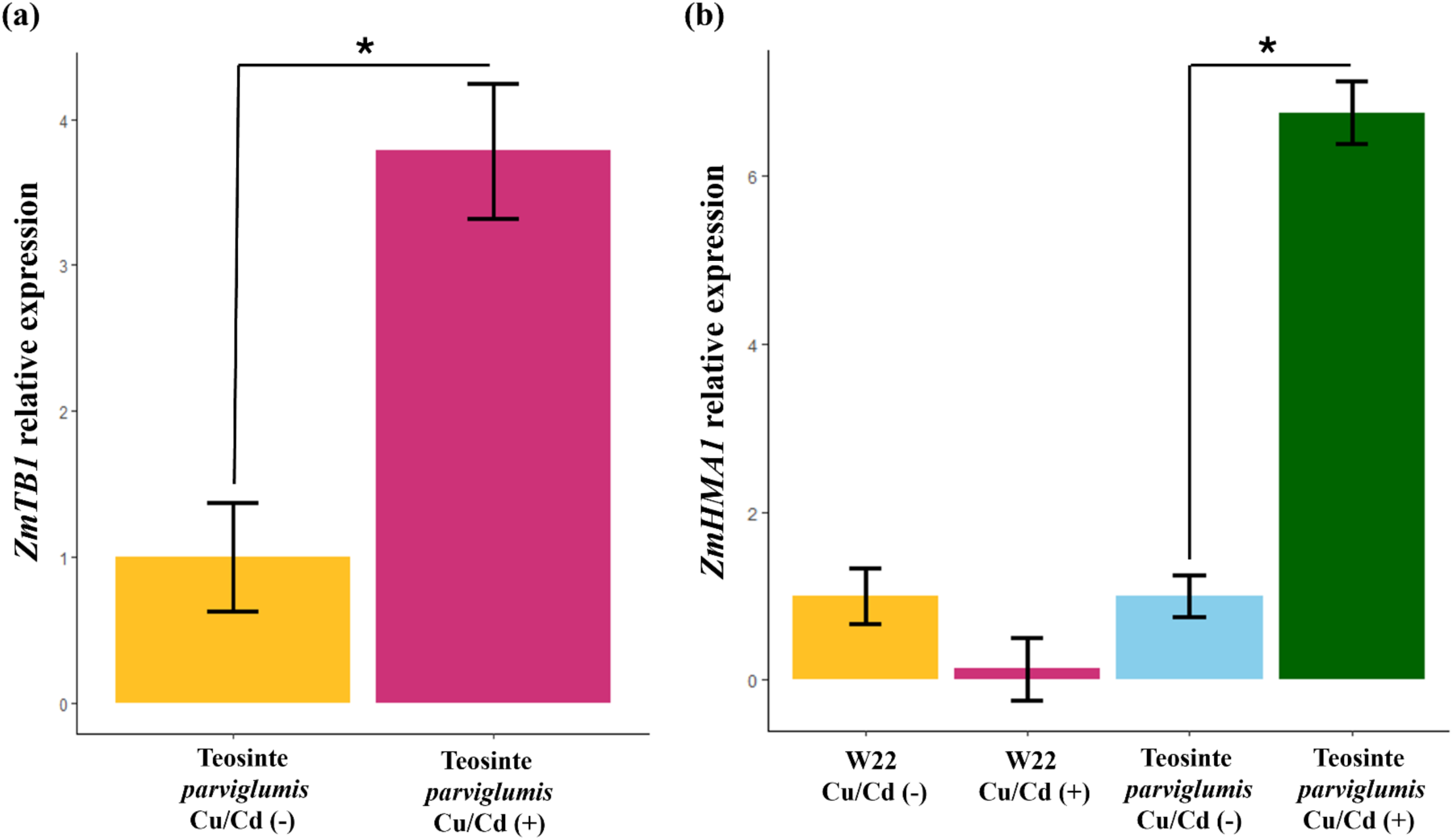
*Tb1* and *ZmHMA1* expression in the shoot apical meristem of plantlets grown under heavy metal stress. **(a**) qPCR analysis of *Tb1* expression in 1-month old teosinte *parviglumis* plantlets grown in absence or presence of heavy metal stress. (**b**) qPCR analysis of *ZmHMA1* expression in 1-month old teosinte *parviglumis* and W22 maize plantlets in absence or presence of heavy metal stress. Histograms show the relative fold-change (normalized to actin, based on technical and biological triplicates); standard deviations are shown by vertical lines and Student T-test P-values represented by asterisks; * P<0.01.

### Paleoenvironmental studies reveal periods of climatic instability in the presumed region of maize emergence during the early Holocene

It is well accepted that temperature fluctuations, volcanism and anthropogenic impact shaped the distribution and abundance of plant species in the Transmexican Volcanic Belt (TMVB) during the last 14,000 years (Torrescano-Valle et al. 2019). The TMVB has produced close to 8000 volcanic structures (Ferrari et al., 2011), transforming the relief multiple times, and causing hydrographic and soil changes that actively modified the distribution and composition of plant communities in Central Mexico. Detailed paleoenvironmental data for the Pleistocene and Holocene is available for several lacustrine zones located within the 50 to 100 km range of the region currently considered the cradle of maize domestication (Matzuoka et al. 2002; Figure 5a). In Lake Zirahuén (102°44′ W; 19°26′ N and approximately 2075 meters above sea level; index [i] in Figure 5a), pollen, microcharcoal and magnetic susceptibility analyses of two sedimentary sequences reveals three periods of major ecological change during the early and middle Holocene. Between 9500 and 9000 calibrated years before present (cal yr BP), pine forests seem to have been associated with summer insolation increases. A second peak of forest change occurred at around 8200 cal yr BP, coinciding with cold oscillations documented in the North Atlantic. Finally, events occurred between 7500 and 7100 cal yr BP shows an abrupt change in the plant community related to humid Holocene climates and a presumed volcanic event (Lozano-García et al., 2013). The environmental history of the central Balsas watershed has also been documented by pollen, charcoal, and sedimentary analysis conducted in three lakes and a swamp of the Iguala valley (Piperno et al. 2007). Paleoecological records of lake Ixtacyola (8°20N, 99°35W and approximately 720 meters above sea level; index [ii] in Figure 5a) and lake Ixtapa (8°21N, 99°26W) indicate that an important increase in temperature and precipitation occurred between 13000 and 10000 cal yr BP. The pollen record of Ixtacyola showed that members of the genus *Zea* were already part of the vegetation coverage by 12900 to 13000 cal yr BP, suggesting that some teosintes – likely including *parviglumis* - were commonly found at elevation areas where they do not presently occur. Lake Almoloya (also named Chignahuapan; 19°05N, 99°20E and approximately 2575 meters above sea level; index [iii] in Figure 5a) in the upper Lerma basin is only 20 Km from the crater of the Nevado de Toluca that is responsible for creating the late Pleistocene Upper Toluca Pumice layer over which the Lerma basin is deposited. Pollen records indicate the presence of *Zea* species by 11080 to 10780 cal yr BP. As for other locations, an important period of climatic instability prevailed between 11500 and 8500 cal yr BP (Ludlow-Wiechers et al., 2005). Humidity fluctuations occurred until 8000 cal yr BP, with a stable temperate climate between 8500 and 5000 cal yr BP. Although pollen and diatom studies are often difficult to interpret at a regional scale, the overall reported results suggest consistent periods of *Zea* plants present in periods of environmental and climatic instability that correlate with the history of volcanic activity during the early Holocene, as described in the next section.

**Figure 5.**
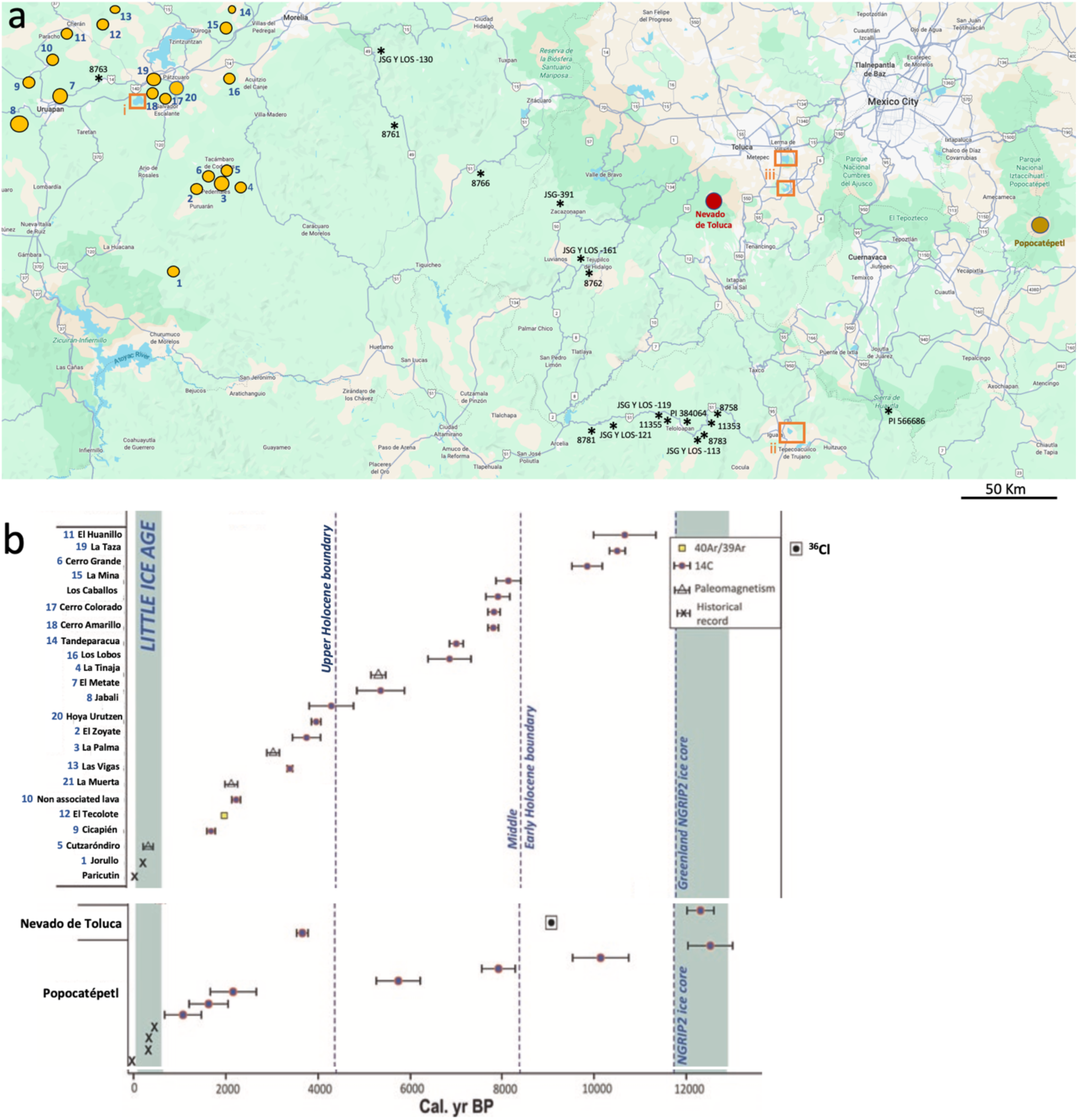
Location and age of volcanic eruptions occurred during the Holocene in comparison to collection sites of teosinte *parviglumis* populations most closely related to maize. (**a**) Black asterisks correspond to the location and accession numbers of collected teosinte *parviglumis* populations that are phylogenetically most closely allied with extant maize, following Matzuoka et al. 2002. The location of 21 volcanoes belonging to the Michoacan-Guanajuato Volcanic Field is indicated with yellow circles. The locations of the Nevado de Toluca and Popocatéptl craters are indicated with a red and brown circle, respectively. Orange squares indicate the location of paleoenvironmental studies covering from 14,000 cal yr BP to current era, as described in Lozano-García et al. 2013 (i), Piperno et al. 2007 (ii), and Ludlow-Wiechers et al. 2005 (iii). (**b**) Time versus age of volcanic eruptions occurred in the map area of the Transmexican Volcanic Belt during the Holocene; numbers correspond to volcanoes illustrated in (**a**). Adapted from Macías and Arce (2019).

### Temporal and geographical convergence between volcanic eruptions and maize emergence during the Holocene

Current evidence indicates that the emergence and domestication of maize initiated in Mesoamerica some time around 9,000 yr BP (Matsuoka et al. 2002). The current location of teosinte *parviglumis* populations that are phylogenetically most closely allied with maize are currently distributed in a region located between the Michoacan-Guanajuato Volcanic Field (MGVF) at their northwest, and the Nevado de Toluca and Popocatéptl volcanoes at their east and northeast (Figure 5a; Matsuoka et al. 2002). Precise records of field data indicate that ten accessions were collected in the Balsas river drainage near Teloloapan and Sierra de Huautla (Guerrero), at approximately 100 km south of the Nevado de Toluca crater (Matzuoka et al. 2002). Three other accessions were collected near Tejupilco de Hidalgo and Zacazonapan (Estado de México), at approximately 50 to 60 km from the Nevado de Toluca crater (8762, JSG y LOS-161, and JSG-391). And four other accessions were located in Michoacan, at a location within the MGVF (accession 8763), or at mid-distance between the MGVF and the Nevado de Toluca crater (accessions JSG y LOS-130, 8761, and 8766).

The most important source of HMs in ancient soils of Mesoamerica is TMBV-dependent volcanic activity through short- and long-term effects related to lava deposits, ores, hydrothermal flow, and ash (Torrescano-Valle et al. 2019). Interestingly, TMVB activity in the center of Mexico was particularly intense during the Holocene, as compared to other regions and time frames (Marcias and Arce, 2019). The Nevado de Toluca volcano produced one of the most powerful eruptions from central Mesoamerica in the Holocene, giving rise to the Upper Toluca Pumice deposit at 12621 to 12025 cal yr BP (Arce et al., 2003; Figure 5b). The pumice fallout blanketed the Lerma and Mexico basins with 40 cm of coarse ash (Bloomfield and Valastro 1977; Arce et al. 2003). A second eruption dated by ^36^Cl exposure occurred at 9700 cal yr BP (Arce et al. 2003; Figure 5b), and the most recent eruption occurred at 3580 to 3831 cal yr BP (Macías et al. 1997). During the early and middle Holocene, the Popocatéptl volcano produced at least four eruptions dated 13037-12060, 10775–9564, 8328-7591, and 6262-5318 cal yr BP (Siebe et al. 1997); three other important eruptions occurred during the late Holocene, between 2713 and 733 cal yr BP (Siebe and Macías, 2006). In addition, the MGFV is a monogenetic volcanic field for which 23 independent eruptions have been documented during the Holocene, 21 of them located towards the southern part of the field, in close proximity to the region harboring some of the teosinte *parviglumis* populations most closely related to maize. Three of these eruptions occurred in the early Holocene (El Huanillo 1130 to 9688 cal yr BP; La Taza 10649 to 10300 cal yr BP; Cerro Grande 10173 to 9502 cal yr BP; Figure 5b), and three others during the initial period of the middle Holocene, between 8400 and 7696 cal yr BP (La Mina, Los Caballos, and Cerro Amarillo; Figure 5b). On average, a new volcano forms every ∼435 years in the MGFV (Macías and Arce, 2019). No less than 16 other eruptions occurred between 7159 cal yr BP and the present time (Figure 5b). Soils of volcanic origin (andosols) are currently distributed in regions north-west from the Nevado de Toluca and Popocatéptl craters, in close proximity with teosinte *parviglumis* populations most closely related to maize (Figure S5). Although modern distribution of teosinte populations may differ from their distribution around 9000 yr BP, and unknown populations more closely related to maize may yet to be discovered, this data indicates that the date and region where maize emerged is convergent with the dates and locations of several volcanic eruptions occurred during the Holocene in that same region.

## Discussion

Although the evolutionary transition from teosinte *parviglumis* to maize resulted in important phenotypic divergence between landraces and their wild progenitor, the relevance of teosinte *parviglumis* phenotypic plasticity and its genetic consequences during domestication remain poorly understood. Our study offers the first evidence showing that a specific environmental factor (HM stress) influenced the teosinte *parviglumis* to maize transition through specific gene function (HM-response ATPases). The comparison of HM responses between both subspecies indicates that teosinte *parviglumis* phenotypically responds to HM stress acquiring phenotypes reminiscent of maize. Low temperatures and low CO_2_ concentrations caused a significant decrease in lateral branching but not in tillering. This decrease of lateral branching correlated with a decrease in teosinte seed yield (Piperno et al. 2015). Similar results were reported for *parviglumis* plants grown under low Nitrogen conditions (Gaudin et al. 2011). Our results under HM stress also show an increase in the number of female inflorescences per plant. This HM stress-dependent proliferation of several closely associated female inflorescences in a single teosinte *parviglumis* stem has not been previously reported. Female inflorescences arose from individual axillary meristems in multiple internodes, significantly increasing teosinte *parviglumis* seed yield. Under HM stress, we also show that *Tb1* is overexpressed in the apical meristem of teosinte *parviglumis,* suggesting that formation of secondary ramifications is repressed by *Tb1* function under HM stress, as in extant maize. At this stage we cannot discard the possibility that *Tb1* upregulation in *parviglumis* reflects a more generalized response to abiotic stress; however, the expression *ZmHMA1* is downregulated in W22 wild-type maize meristems in the presence of HMs but upregulated in teosinte *parviglumis* meristems, suggesting that a specific regulatory shift relating HM responses and *ZmHMA1* function occurred during the teosinte *parviglumis* to maize transition.

We also show that *ZmHMA1*, *ZmHMA7* and *ZmSKUs5* exhibit reduced genetic diversity in extant maize as compared to teosinte *parviglumis*, suggesting that all three loci were affected by the evolutionary transition that gave rise to maize. The maize regulatory region and downstream 3’ area of the *ZmHMA1* locus exhibit pronounced reduction in SNPs compared to teosinte *parviglumis* lines, likely affecting the molecular scope of its regulatory proteins, including transcription factors of the TCP family (Martín-Trillo and Cubas 2010). By contrast, *ZmHMA7* and *ZmSKUs5* showed a selective sweep across the entire maize locus. Although loss of genetic diversity is usually the result of human selection during domestication, it can also represent a consequence of natural selective pressures favoring fitness of specific teosinte *parviglumis* allelic variants better adapted to environmental changes and subsequently affected by human selection during the domestication process. This possibility is reflected by widely spread selective sweeps affecting a large portion of chr.5 that contains hundreds of genes showing signatures of positive selection. The analysis of 11.47 Mb covering the *ZmHMA1-ZmSKUs5* segment confirms the presence of large but discrete genomic subregions that were positively selected during the teosinte *parviglumis* to maize transition. Although several contain genes involved in HM response and oxidative stress, the diversity of gene functions does not necessarily favor abiotic stress over other factors that could be at the origin of selective forces affecting these regions. By contrast, a large scale transcriptomic survey indicates that genes consistently responding to HMs (Cu, Cd, Pb and Cr ) show signatures of positive selection at unusual high frequencies (43.3%) as compared to loci containing genes responding to other types of abiotic stress (28.6%). Our identification of HM response genes affected by positive selection is far from being exhaustive. Nevertheless, it agrees with the expected effects of a widespread selective sweep caused by environmental changes that influenced the *parviglumis* to maize transition at the genetic level. Of intriguing interest are 24 loci that partially or completely lack SNPs in both teosinte *parviglumis* and maize, suggesting possible genetic bottlenecks occurred before the teosinte to maize transition. Examples of other edaphological factors driving genetic divergence either in the teosintes or maize include local adaptation to phosphorus concentration in *mexicana* and *parviglumis* (Aguirre-Liguori et al. 2019), and fast maize adaptation to changing iron availability through the action of genes involved in its mobilization, uptake, and transport (Benke and Stich 2011). Our results reveal a teosinte *parviglumis* environmental plasticity that could be related to the function of HM response genes positively selected during the teosinte *parviglumis* to maize transition. Previous studies have demonstrated that transposable elements (TEs) contribute to activation of maize genes in response to abiotic stress, affecting up to 20% of the genes upregulated in response to abiotic stress, and as many as 33% of genes that are only expressed in response to stress (Makarevitch et al., 2015). It is therefore possible that the HM response of some specific genes that influenced maize emergence or domestication could be mediated by TEs influencing or driving their transcriptional regulation.

The maize genome contains 12 genes encoding for members of the P1b family of ATPases (Cao et al. 2019). They are subdivided according to their preferential substrate affinity: the Copper (Cu)/Silver (Ag) group includes *ZmHMA1* to *ZmHMA4*; and the Zinc (Zn)/Cobalt (Co)/Cadmium (Cd)/Lead (Pb) includes *ZmHMA5* to *ZmHMA12*. Initial studies suggested that *ZmHMA1* is expressed at low levels in vegetative, reproductive or root organs throughout development, contrary to *ZmHMA7* that is strongly expressed in developing seeds 10 days after pollination; however, both genes appeared downregulated in plants grown under either Cu or Cd stress (Zhiguo et al. 2018). Additional studies by Cao et al (2019) indicated low expression of *ZmHMA1* and high levels of *ZmHMA7* expression in 12 tissues: roots, nodes, shoots, leaves, tassel, shoot apical meristem and young stem, internodes, anthers, silks, whole seedlings, endosperm, embryo, and pericarp. Under Cd stress, expression of both *ZmHMA1* and *ZmHMA7* was significantly upregulated in roots and downregulated in stems (Cao et al. 2019), contradicting previous results by Zhiguo et al. (2018). In Arabidopsis and barley, the corresponding *ZmHMA1* structural homologs (*AtHMA1* and *HvHMA1*, respectively) can transport a wide range of metals into the chloroplast (Mikkelsen et al. 2012; Boutigny et al. 2014). In the case of the Arabidopsis structural homolog of ZmHMA7, AtHMA7 (also called RAN1) is proposed to act as a metallochaperone located in the Golgi apparatus, delivering Cu to the ethylene receptor located in the plasma membrane (Hirayama et al. 1999). Our evidence complements previous studies by showing that in maize, *ZmHMA1* root expression is restricted to vegetative developmental stages, and that *ZmHMA7* is expressed in roots in response to HM stress. We also show through qPCR analysis that under HM stress, the expression of both *ZmHMA1* and *ZmHMA7* is upregulated in teosinte *parviglumis* primary roots of 4-week-old plantlets. Overall, these results confirm that both *ZmHMA1* and *ZmHMA7* are HM response genes active throughout teosinte and maize development.

In maize, the phenotypic analysis of null mutant individuals indicates that *ZmHMA1* is involved in the control of aerial and underground developmental traits, some of which are related to important functional adaptations that distinguish maize from its wild ancestor. In addition to changes in leaf number, leaf length, and number of spikes per male inflorescence, *zmhma1-m1* mutant individuals grown under normal edaphological conditions showed an increase in the number of female inflorescences per plant, a reduction in the length of nodal roots, and an increase in the number of seminal roots per plant. Whereas the promotion of nodal root growth is a distinctive trait of maize phenotypic plasticity and adaptation, previous studies showed that contrary to maize, plants of teosinte *parviglumis* lack seminal roots. This absence is also characteristic of paleobotanical maize specimens dating more than 5,000 cal yr BP, confirming that the acquisition of seminal roots is an important trait acquired during domestication (Lopez-Valdivia et al. 2022). The presence of seminal roots distinguishes extant maize from the teosintes (Burton et al. 2013), suggesting a possible involvement in domestication (Perkins and Lynch 2021; Lopez-Valdivia et al. 2022). Our results suggest that the allelic variants of the *ZmHMA1* locus selected during domestication could have contributed to promote seminal root formation in maize, optimizing their number in accordance to seed endosperm availability and carbohydrate reserves (Perkins and Lynch 2021; Lopez-Valdivia et al. 2022). Interestingly, *ZmHMA1* plays an important role in contributing to restricting plant height in response to HM stress, a phenotype that is also an important distinction between teosinte *parviglumis* and maize. Taken together, these results indicate that *ZmHMA1* function was involved in the phenotypic transition from teosinte *parviglumis* to maize through the influence of HM stress; however, a better definition of its role in the origin and domestication of maize will require a functional analysis of the *ZmHMA1* teosinte *parviglumis* homolog, and a detailed molecular analysis of the corresponding allelic variants and their phenotypic effects in both subspecies.

### Ideas and Speculation

Our results point to an evolutionary model in which environmental changes in the form of edaphological HM stress imposed natural selective forces that acted on native teosinte *parviglumis* populations, causing phenotypic responses and a severe genetic bottleneck (Figure S6). Adapted plants developed phenotypic traits such as shorter height and emergence of seminal roots, progressively giving raise to primitive maize individuals with reduced genetic variation in HM response genes such as *ZmHMA1*. The subsequent action of human selection caused a drastic reduction in allelic variation of HM response genes, and genetic fixation of the resulting domestication traits.

The history of eruptive activity in the TMVB during the early Holocene coincides with the temporal estimation of initial stages in the evolutionary transition that gave rise to maize. Intense and recurrent eruptions occurred 12,000 to 6,000 years ago near the region presumed to be the cradle of maize domestication (Matsuoka et al. 2002; Macias and Arce, 2019). Located less than 60 Km from the region that harbors some of the natural teosinte *parviglumis* populations most closely related to maize (Matsuoka et al. 2002), the Nevado de Toluca volcano underwent three major eruptions approximately 12621-12025, 9700, and 3580- 3831 yr BP (Macias and Arce, 2019). The large 9700 yr BP eruption caused in the region a period of climatic instability that lasted until 8500 yr BP (Ludlow-Wiechers et al., 2005). Could volcanic eruptions during the early Holocene have modified soil composition and influence the initial evolutionary transition from teosinte *parviglumis* to primitive maize? The possibility of environmental cues influencing maize emergence was mentioned by Benz and Iltis (1992), but not in relation to volcanic activity. Deciphering the ancient DNA sequence of HM response genes such as *ZmHMA1* in paleobotanical maize specimens found in Central Mexico could help determine if its genetic diversity was already reduced by 5,300 to 6,200 yr BP (Ramos-Madrigal et al., 2016 ; Vallebueno-Estrada et al., 2016; Benz, 2001). Our results open the possibility for environmental changes caused by volcanic activity acting as a driving force in the selection of loci involved in adaptive responses such as heavy metal tolerance.

## Materials and Methods

### Plant materials and growth conditions

Seeds from maize inbred line W22, teosinte *parviglumis* accession CIMMYTMA 29791, and insertional line *LANMu1029790* were sterilized using a 1:1 distilled water and bleach solution, rinsed three times in water, and germinated in MS solid medium. One week-old seedings were transplanted into PVC tubes (length: 90 cm; diameter: 20 cm) or 20-liter pots filled with a pre-sterilized mix composed of 30% peat moss, 10% vermiculite, 10% perlite, and 50% ground litter, complemented with 15-9-12 (N-P-K) Osmocote^TM^ fertilizer.

For HM growth conditions, 16 mg/kg of cadmium chloride (CdCl_2_) and 400 mg/kg of copper sulfate (CuSO_4_) were added to the substrate before seedling transplant. All plants were grown under glasshouse conditions at 28°C during day and 24°C during night, with relative humidity comprised between 40 and 50%.

### Phenotypic analysis

All plants were harvested at 15 days (V2 stage), 30 days (V6 stage), and 50 days (R1 stage) after transplant. No less than ten plants per biological replicate were measured for each trait and stage, with three biological replicates per experiment. The width and length of immature leaves was measured in the 5^th^ leaf. The length and width of mature leaves in the 3^rd^ leaf. The total number of leaves, total number of stem internodes, plant height, basal stem diameter, total number of stem ramifications, and total fresh and dry weight of the aerial biomass were estimated at the R1 stage. Estimated root traits included the number and length of brace roots (aerial roots), the number and length of crown roots (underground roots originating within 2 cm below the basal node), number and length of nodal roots (underground roots originating more than 2 cm below the basal node), the number and length of seminal roots and maximum length of the root system. Statistical analyses were conducted using RStudio, version 4.3.0 (Horton and Kleinman 2015). The data are presented as mean ± standard deviation and were tested by a paired Student’s T-Test. When the effects were significant according to T-Test, the treatments were compared with the Welch two sample T-Test at *P <* 0.05.

### Genetic diversity analysis

The SNP collection of maize chromosome 5 was analyzed on the basis of the HapMap V3.2.1 dataset <u>(Bukowski et al. 2018)</u>, selecting individuals from maize landraces (LR) (Table S6) and teosinte *parviglumis* inbred line accessions (Teo; Table S7). A sliding window diversity (π) was calculated using VCFtools v0.1.13 <u>(Danecek et al. 2011)</u> with a window size of 100 bp (--window-pi 100) and a sliding step of 50 bp (--window-pi-step 50). Regions under selection were extracted from a list of 304 genes with domestication-related selection signatures across the whole chromosome (Hufford et al. 2012; Tables S8), for a total of 385,829 sliding windows (Table S9 and S10). A second set of neutral gene regions comprising 2,373,010 windows was also extracted across the entirety of chromosome 5. These two collections were compared to *ZmHMA1* (GRMZM2G067853), *ZmHMA7* (GRMZM2G029951), and *ZmSKUs5* (GRMZM2G076985), encompassing 2,743 windows (Table S11 and S12). Additionally, we analyzed genetic diversity in the neighboring regions of these candidate genes, which included six genes (*GRMZM2G701784*, *GRMZM2G014508*, *GRMZM2G367857*, *GRMZM2G375607*, *GRMZM2G374375*, and *GRMZM2G067883*), totaling 3,088 windows. We employed a custom Perl script (<u>Vallebueno GitHub repository</u>) to calculate two diversity metrics: fold change (FC= π(LR)/π(Teo)) and the proportional diversity (FP=π(LR)/(π(LR) + π(Teo)). Data visualization was performed using violin plots generated in R v4.2.2.

For an arbitrary locus encompassing 30 Kbs up and downstream of the coding sequence of *ZmMA1*, *ZmHMA7*, and *ZmSKUs5*, the number of segregating sites (S), the number of unique sequences (haplotypes, h), and the nucleotide diversity index (θ) were estimated using DnaSP version 6.0 for all accessions included in HapMap3 (Rozas et al. 2017; Bukowski et al. 2018). DnaSP was also used to estimate Tajima’s D, Fu and Li’s D, and Fu and Li’s F indexes (Tajima 1989; Fu and Li 1993). The average proportion of pairwise nucleotide differences per nucleotide site (π) was calculated with VCFtools using bins of 100 bp and steps of 25 bp (Tajima 1983; Weir and Cockerham 1984). Null hypothesis of equality of parameters followed two-tailed tests as previously described (Rozas et al. 2017).

To characterize genetic diversity across a 11.47 Mb genomic interval encompassing the ZmHMA1 to *ZmSKUs5* segment on chr.5, we quantified nucleotide diversity (π) differences between maize and teosinte *parviglumis*. For each gene found within the segment, three regions were defined: an upstream region, the coding sequence (CDS), and a downstream region. Nucleotide diversity for each subspecies was calculated within each region and expressed as the ratio πMaize/πTeosinte (πM/πT). Subregions in which maize retained less than 70% of teosinte *parviglumis* diversity (π_M/π_T ≈ 0.7) were highlighted in Supplemental File 6. Genome-wide π estimates were derived from non-overlapping 100 bp windows that had been intersected between species. Two complementary statistical approaches were used to evaluate species differences per region. First, a parametric analysis of variance (ANOVA) was fitted for each region using π as the response variable and subspecies (maize vs teosinte *parviglumis*) as a fixed factor. ANOVA was used when its assumptions (approximate normality of residuals and homogeneity of variances) were considered acceptable following Fischer. Second, a paired Wilcoxon signed-rank test was applied to the paired π values from maize and teosinte *parviglumis* within each region as a distribution-free alternative robust to departures from normality and to outliers (Wilcoxon, 1945). P-values from both methods were adjusted to control the false discovery rate (FDR) using the Benjamini–Hochberg procedure (Benjamini & Hochberg, 1995). All data processing and statistical analyses were performed in R (version 4.3.1). Gene annotations were obtained from Phytozome using the *Zea mays* PHJ40 v1.2 annotation (Bornowski et al., 2021).

### Gene expression analysis

Leaves, stems, tassels, female inflorescences, as well as primary, nodal, and crown roots were manually isolated, frozen in liquid nitrogen and stored at 4°C. Total RNA was extracted using TRIZOL (Thermo Fisher Scientific, USA). For reverse-transcriptase PCR (RT-PCR), RNA was converted to coding DNA using the One-step RT-PCR kit (QIAGEN, USA). The full list of primers is presented in Table S8. *ZmCDK* was used as a positive control. PCR amplification was conducted using the following parameters: at 94°C for 5 min.; 94°C 30 sec., 60°C 30 sec., 72°C 1 min. for 35 cycles; final stop after 72°C for 10 min. For quantitative real-time PCR (qPCR), total RNA was isolated from 4-week-old meristems using TRIZOL. Complementary DNA (cDNA) was synthesized using 4 μg of total RNA, 10 mM oligodTs and Superscript reverse transcriptase II (Invitrogen, USA). Primers for PCR were designed using the online program Primer 3 Plus (v.0.4.0) and verified with Oligocalculator (Sigma Chemical, St Louis MO) to discard dimer formation. PCR efficiencies of target and reference gene were determined by generating standard curves based on serial dilutions prepared for cDNA templates. PCR efficiency was calculated according to the slope of the standard curve (primers with 100% efficiency, the fold equals 2). Each quantitative real-time PCR (qPCR) reaction was performed in a 10 μl volume consisting of 5 μl of 23 SYBR Green PCR reaction Mix (Applied Biosystems, Foster City, CA). 3.5 μl of the DNA template (100 ng/ml), 0.5 μl of the forward primer (5 mM), 0.5 μl of the reverse primer (5 mM), and 0.5 μl of ultrapure water. The qPCR reactions were performed using the CFX96 Touch Real-Time PCR detection System and the data were analyzed using the Bio-Rad CFX Manager software v3.1. The thermal profile consisted of 10 min at 95°C, 40 cycles of 15 sec at 95°C, and 1 min at 60°C. Amplification results were collected at the end of the extension step. Primer sequences for qPCR amplification are listed in Table S7. A comparative 2-ΔΔCT method was used for determining a relative target quantity of gene expression (Schefe et al. 2006), and the cyclin-dependent kinase gene *ZmCDK* (GRMZM2G149286) was used as a control (Hu et al. 2019). Reproducibility of the results was evaluated for each sample by running three technical and three biological replicates of each of the reactions and each sample.

## Acknowledgments

We would like to thank Ueli Grossniklaus for critical comments of this manuscript; Juana de la Cruz for supporting greenhouse activities; Ivan López-Valdivia, Eduardo González-Orozco and Cristian Eduardo Martínez Guerrero for help with bioinformatic analysis; and Enrique Pola for help with qPCR analysis. All members of the Apolab offered valuable suggestions and support. This research was funded by Conahcyt grant (CB-256826) and the INAH-Cinvestav Tehuacán Collaborative Initiative. J.A-B. is the recipient of a graduate scholarship from Consejo Nacional de Humanidades Ciencia y Tecnologia (CONAHCyT).

## Competing interests

None declared

## Author contributions

J. A-B and J.P.V-C designed the study. J. A-B and J.P.V-C coordinated and managed the project. M.V.E. designed and performed computational analysis. J.A-B performed the experiments, J. A-B, M.V.E and J.P.V-C generated, analyzed and interpreted data J. A-B and J.P.V-C wrote the article. All authors read and approved the final version of the manuscript.

**Supplementary File 1**. List of domestication genes present on chr5 as defined by Hufford et al. 2012

**Supplementary File 2**. FC and FP values for all domestication genes present in chr.5.

**Supplementary File 3**. FC and FP values of neutral regions included in chr.5.

**Supplementary File 4**. FC and FP values of candidate genes *ZmHMA1*, *ZmHMA7* and *ZmSKUs5*.

**Supplementary File 5**. FC and FP values of *ZmHMA1*, *ZmHMA7* and *ZmSKUs5* neighboring genes

**Supplementary File 6.** Genetic diversity in loci across 11.47 Mb of chr.5 in the *ZmSKUs5* to *ZmHMA1* segment

**Supplementary File 7.** Genetic diversity in loci encompassing heavy metal response genes shared by six transcriptomes with fold change strictly higher than 1 and FDR<0.05

---

**Figure S1.**
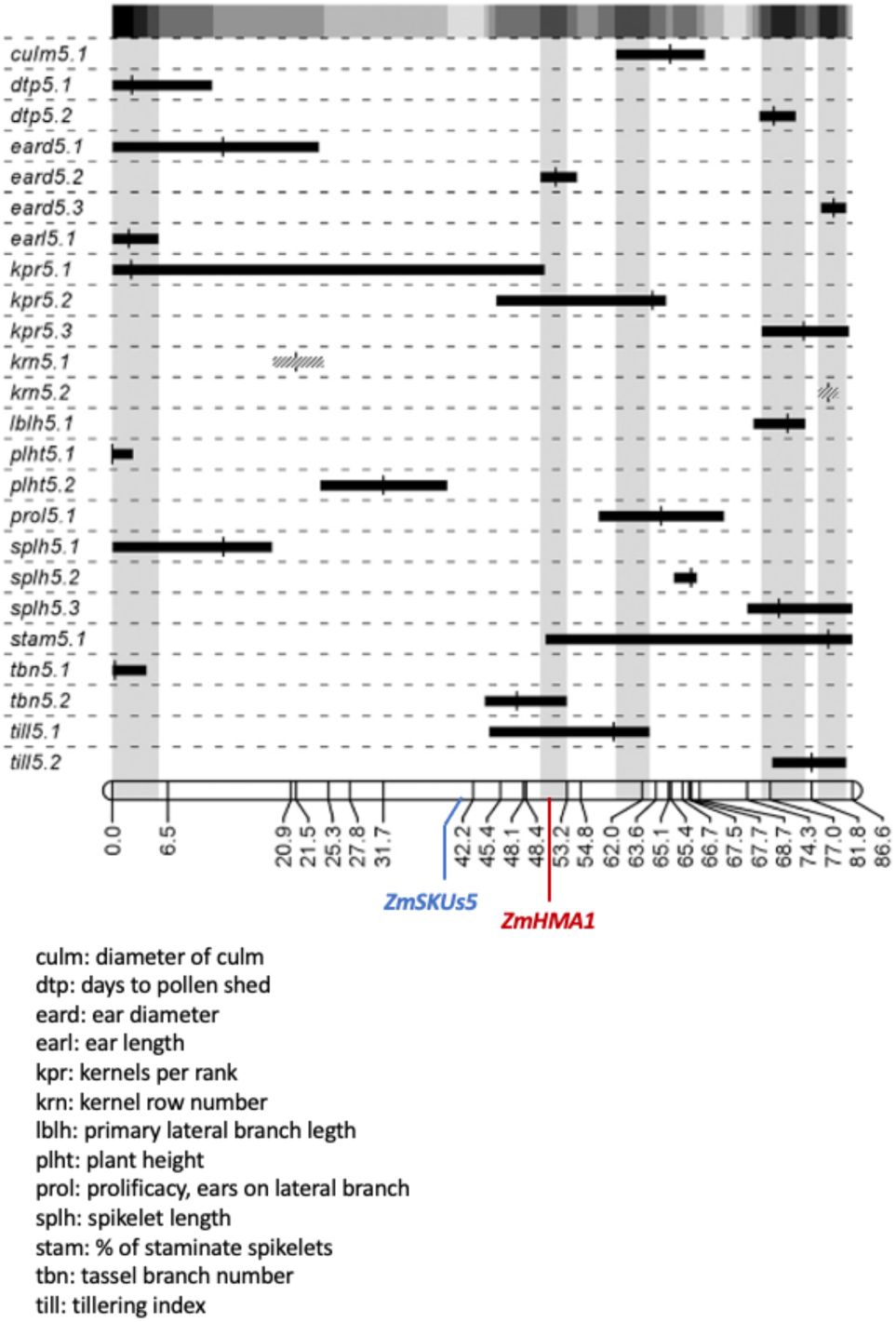
Cumulative plot of QTL detected in a region of chromosome five that includes *ZmSKUs5* and *ZmHMA1* (modified from Lemmon and Doebley, 2014).

**Figure S2.**
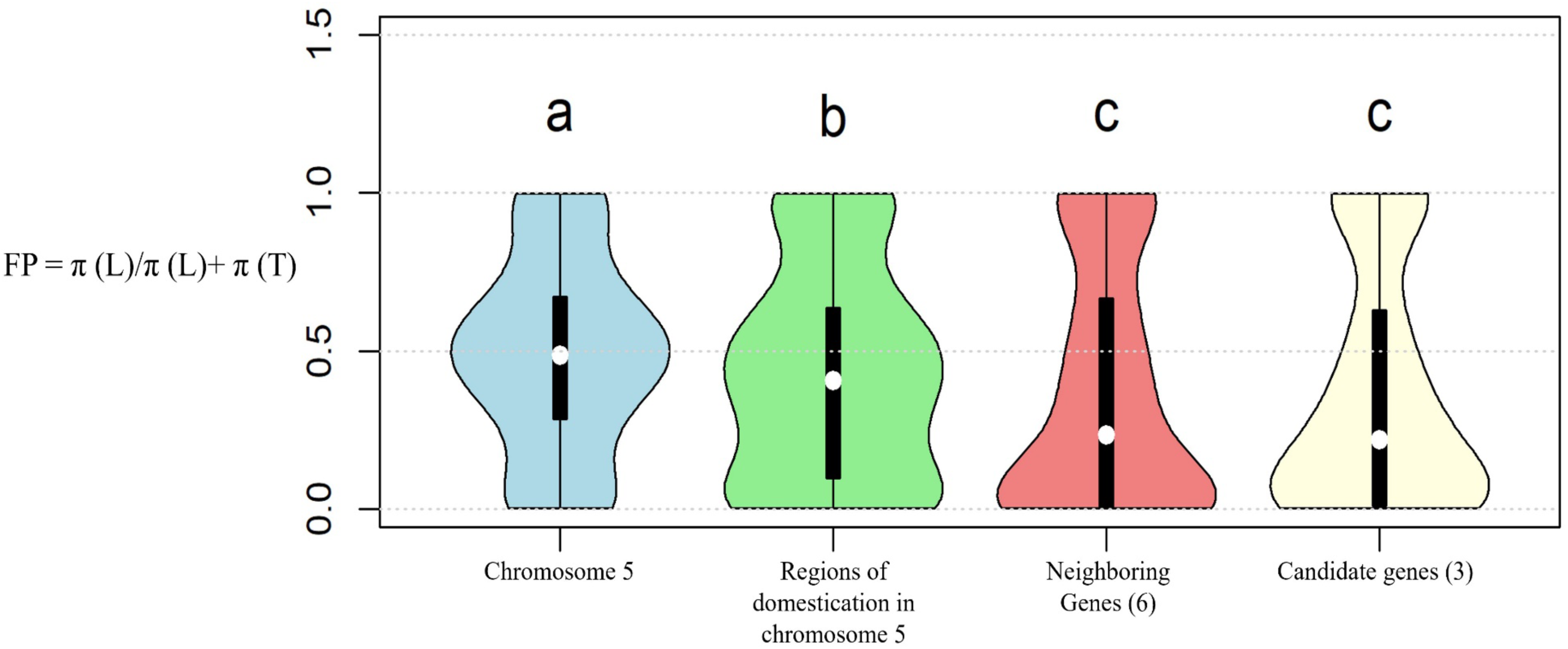
Comparison of chromosome 5 nucleotide variability between maize landraces and teosinte *parviglumis*.

**Figure S3.**
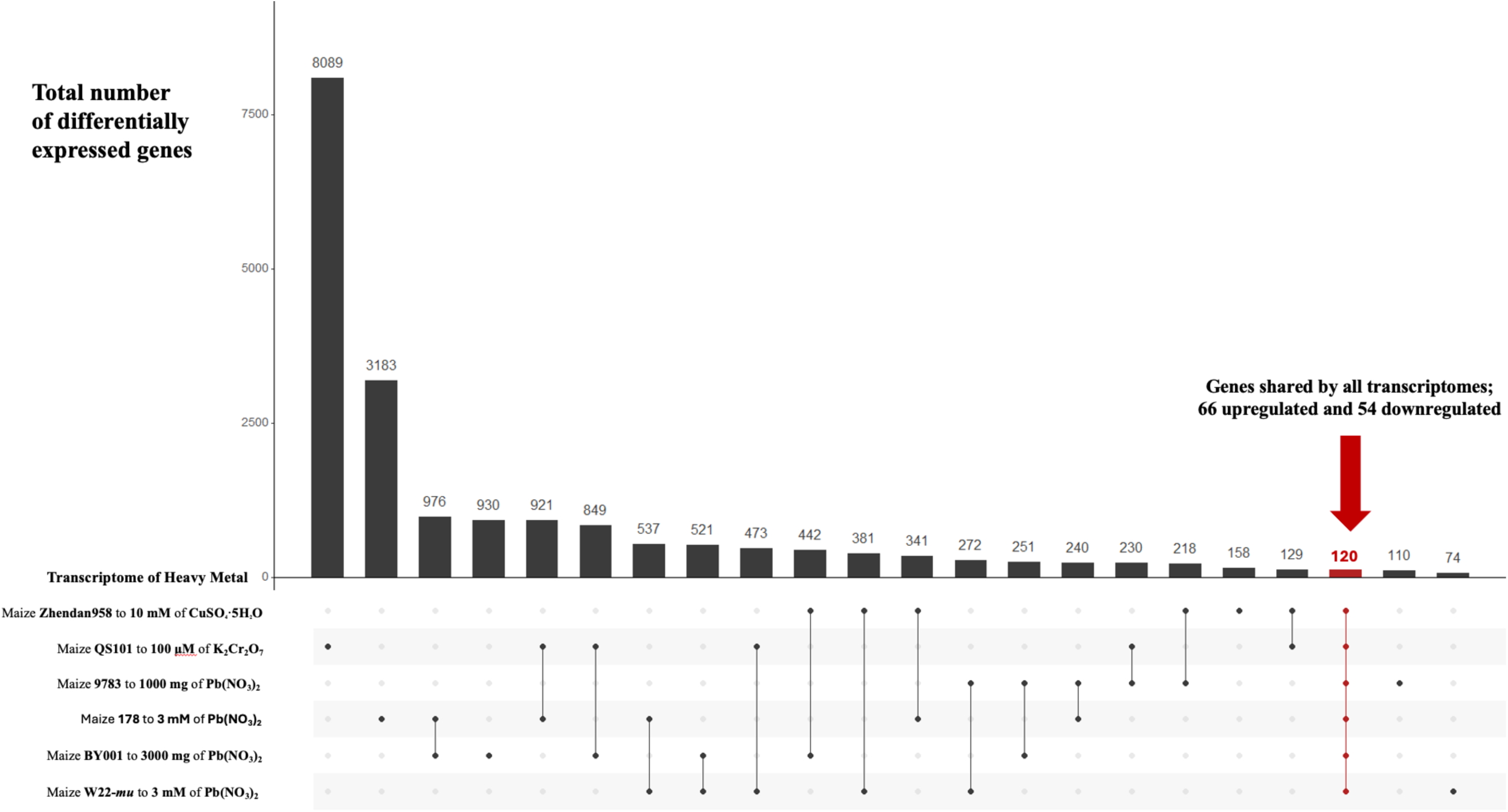
Large scale transcriptional comparison of HM response genes across six maize experiments based on fold change >1 and false discovery rate (FDR) < 0.05.

**Figure S4.**
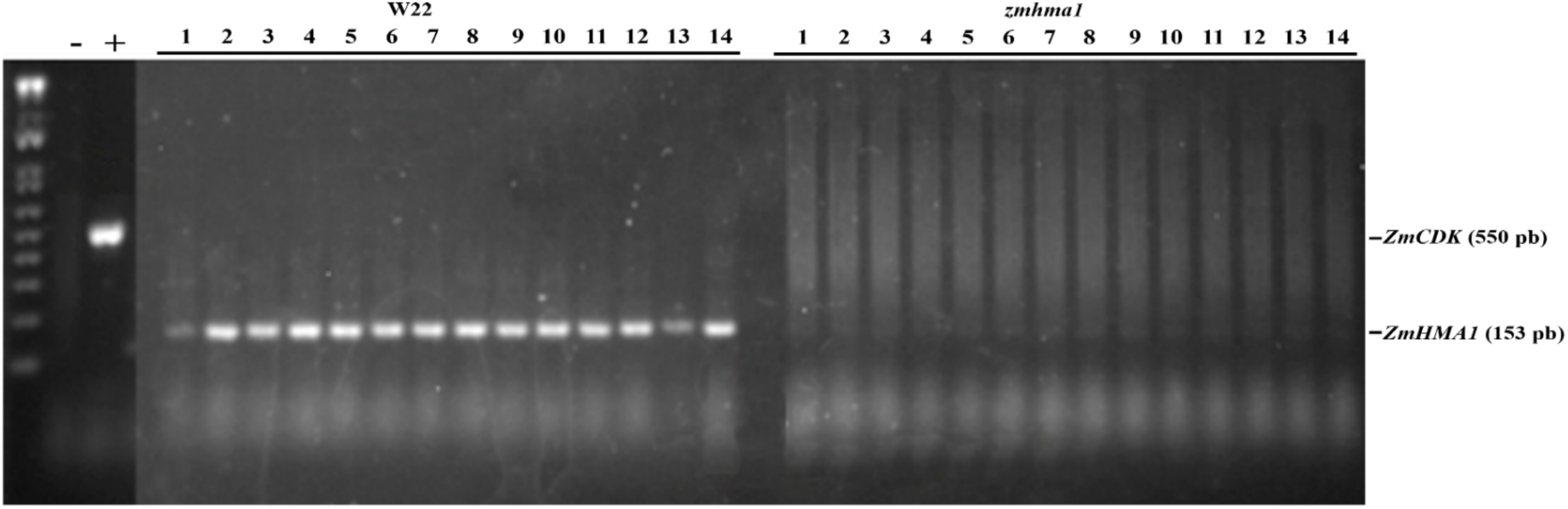
Expression of *ZmHMA1* in wild-type and homozygous *zmhma1* maize individuals.

**Figure S5.**
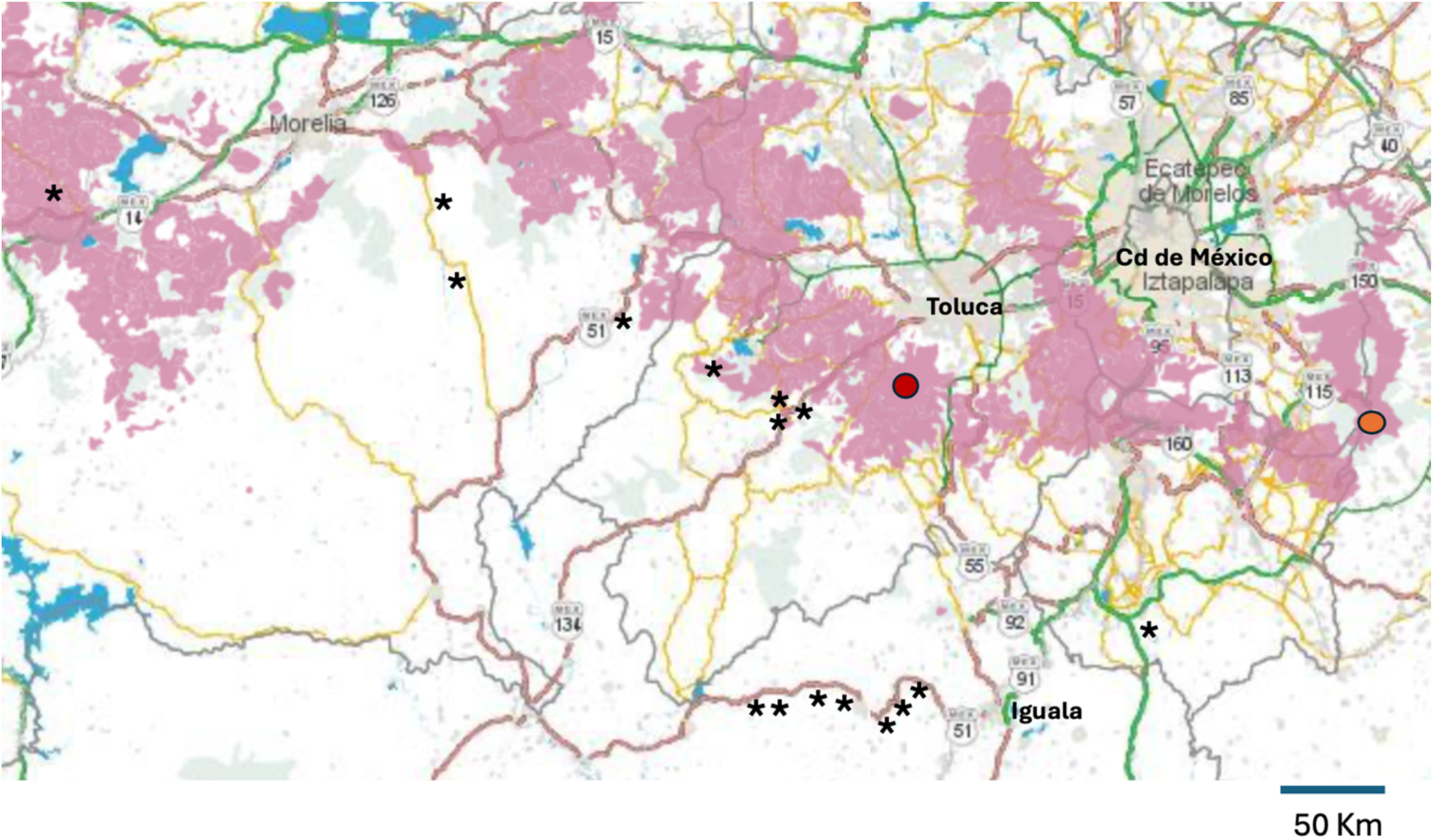
Distribution of soils of volcanic origin (andosols; pink) in geographic regions of teosinte *parviglumis* populations most closely related to extant maize.

**Figure S6.**
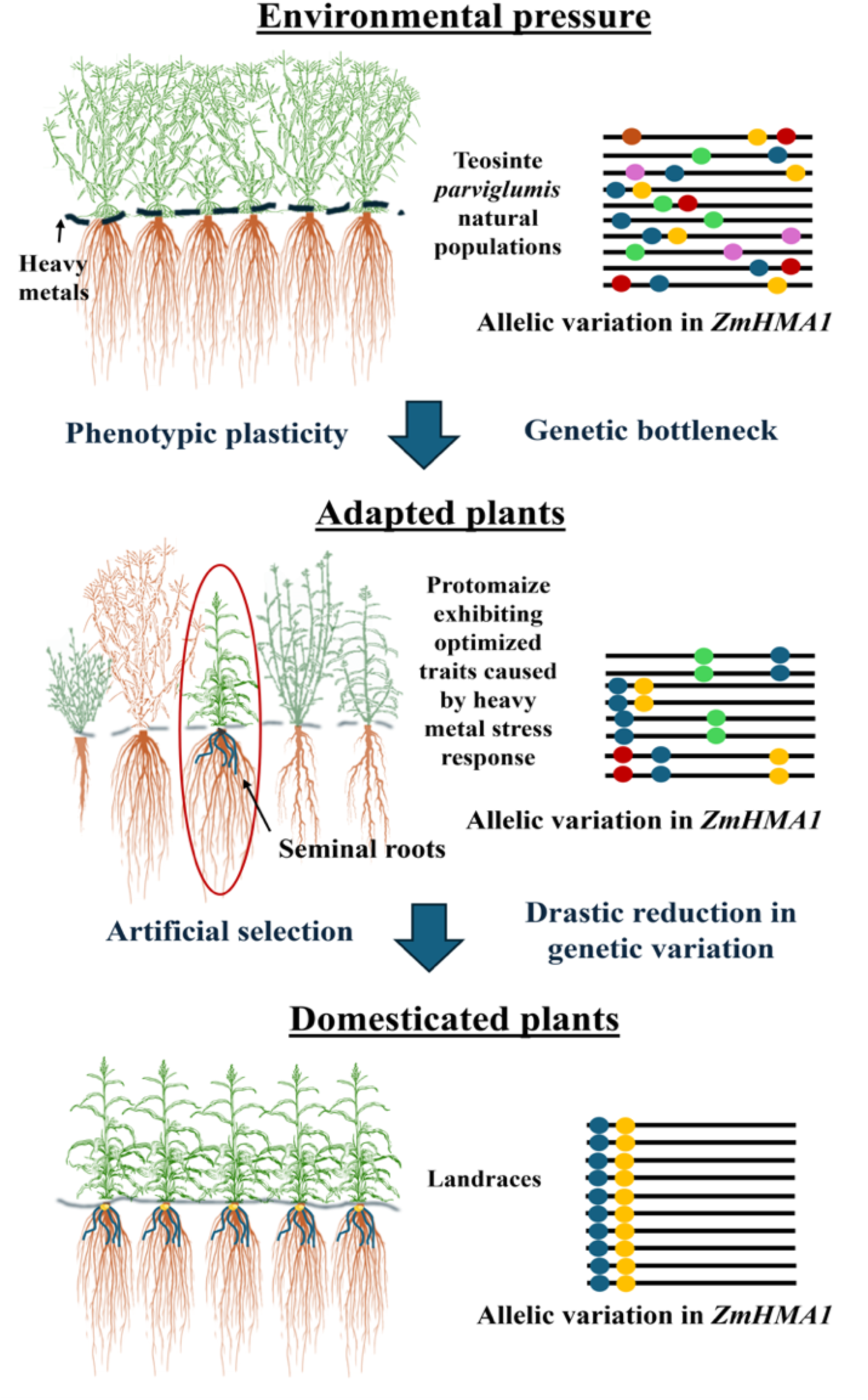
Speculative model illustrating the influence of heavy metal stress on the evolutionary transition of teosinte *parviglumis* to maize.

**Table S1.**
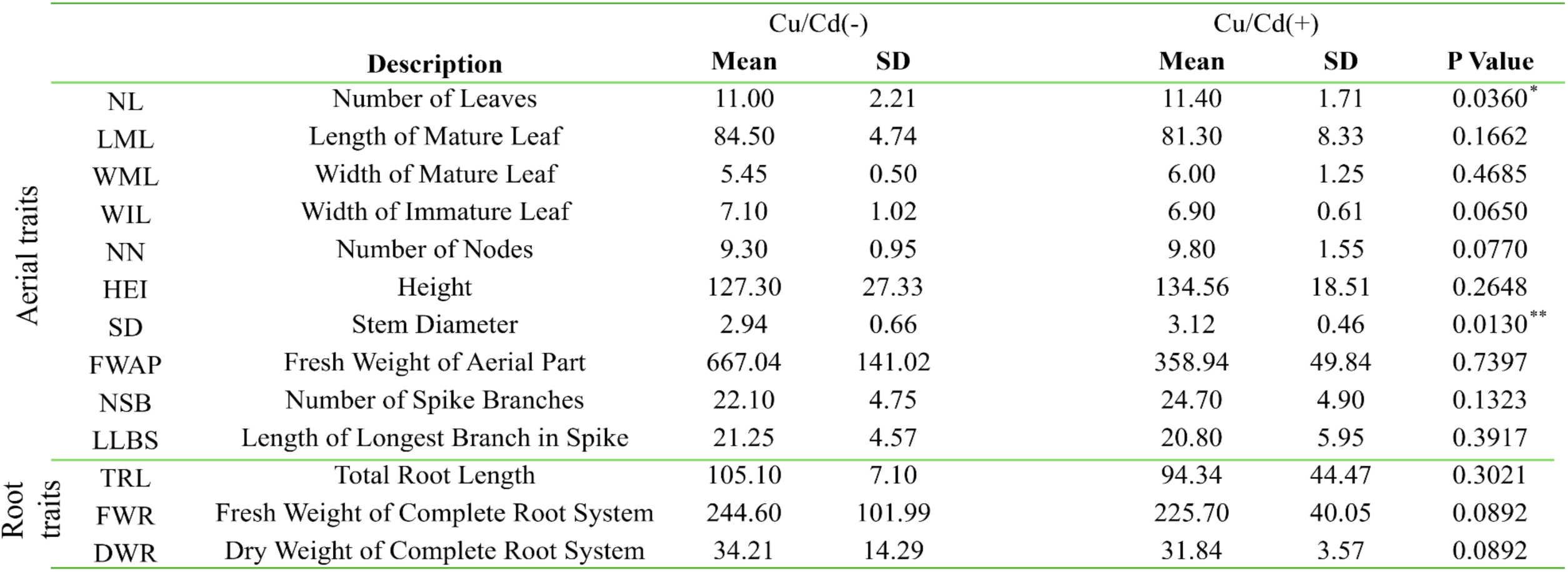
Additional estimation of phenotypic traits in teosinte *parviglumis* grown in absence or presence of heavy metal stress.

**Table S2.**
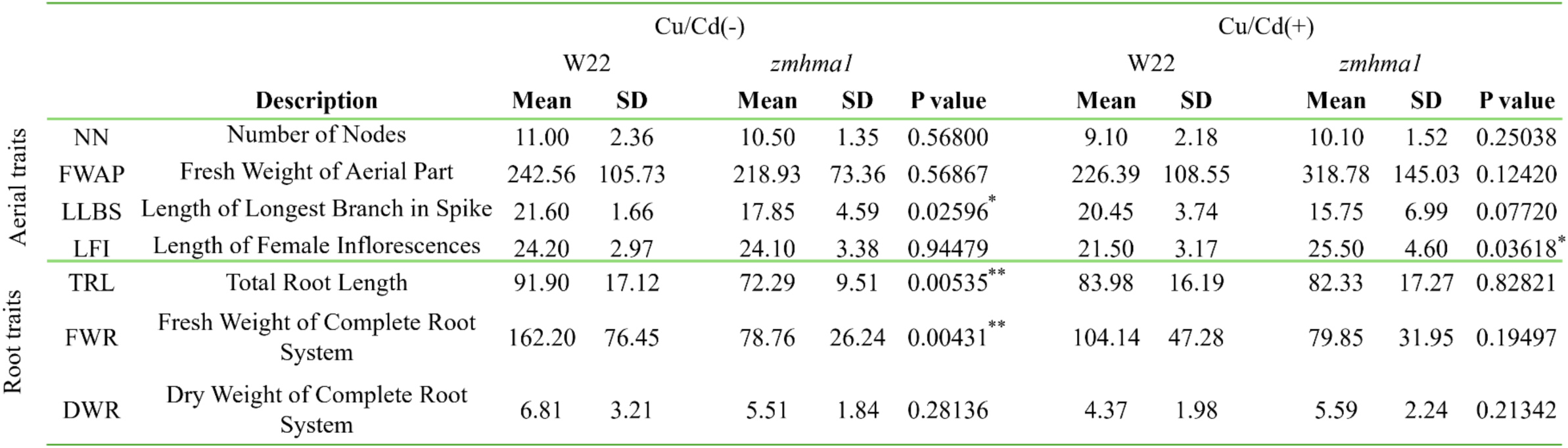
Additional estimation of phenotypic traits in wild-type and *zmhma1* maize grown in absence or presence of heavy metal stress.

**Table S3.**
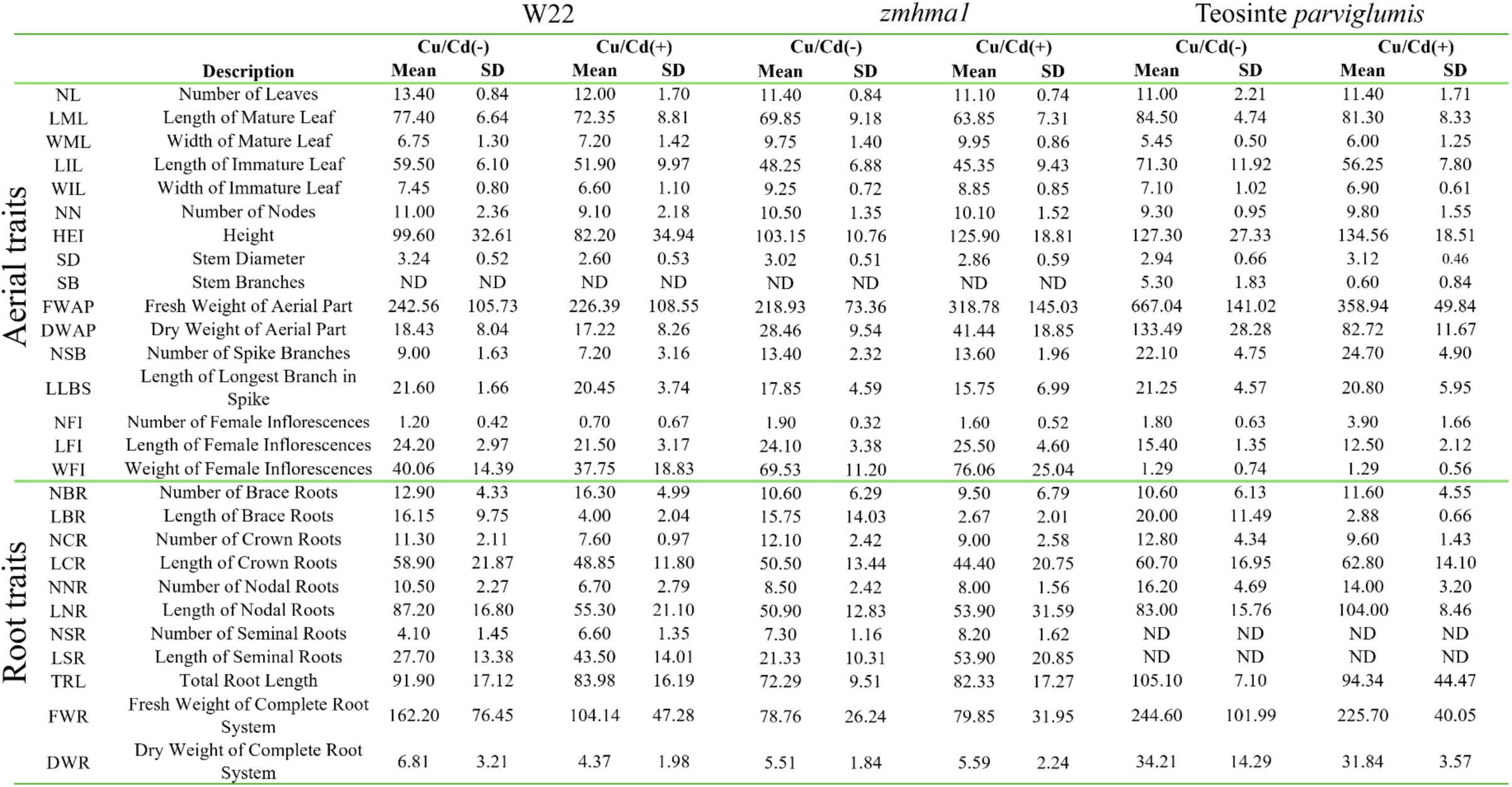
Full comparison of phenotypic values for wild-type W22, *zmhma1*, and teosinte *parviglumis* individuals grown under absence or presence of heavy metal stress.

**Table S4.**
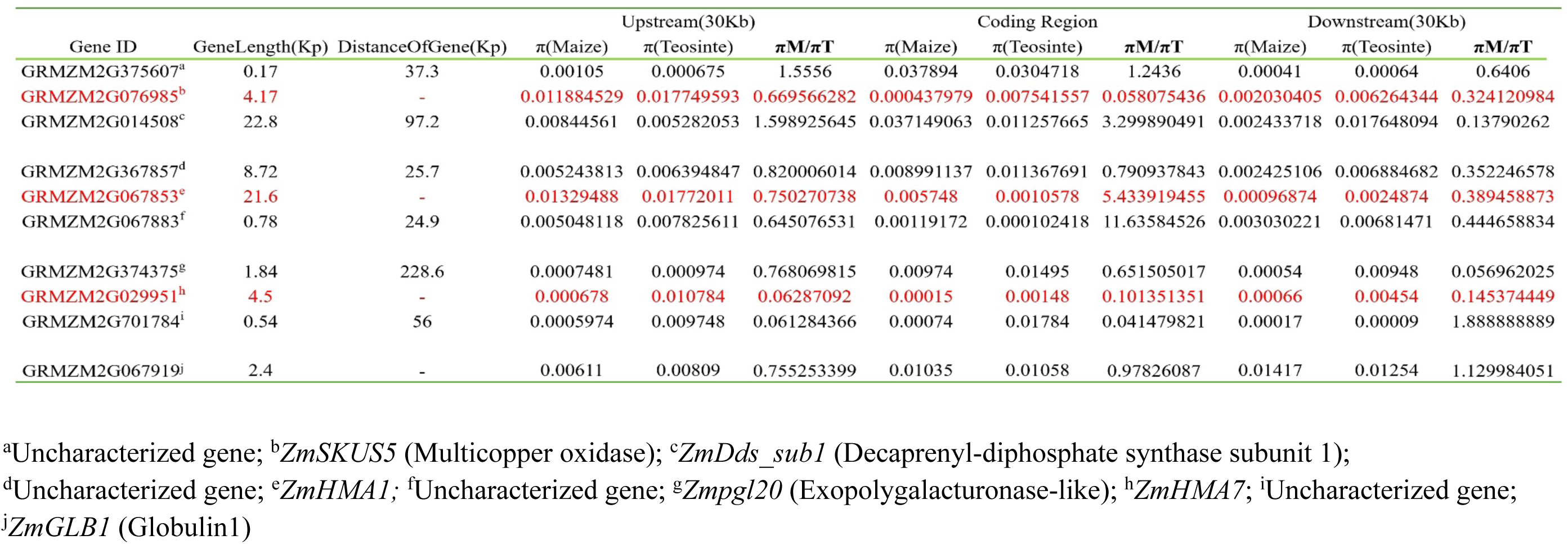
Nucleotide variability across the *ZmSKUS5, ZmHMA*1 and *ZmHMA7* locus; and their upstream and downstream neighboring genes in maize landraces, maize improved lines, and teosinte *parviglumi*s accessions included in HapMap3.

**Table S5.**
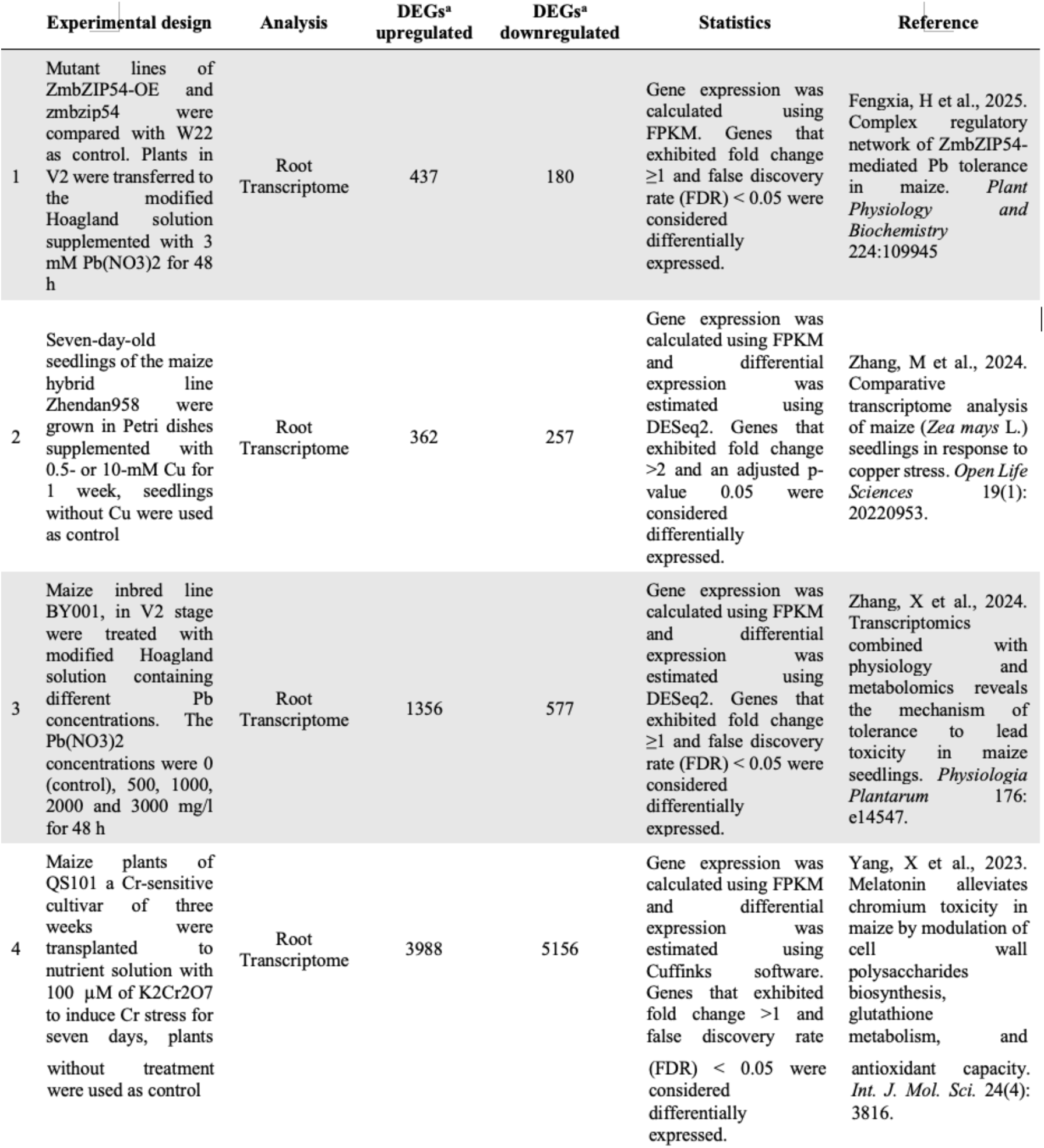
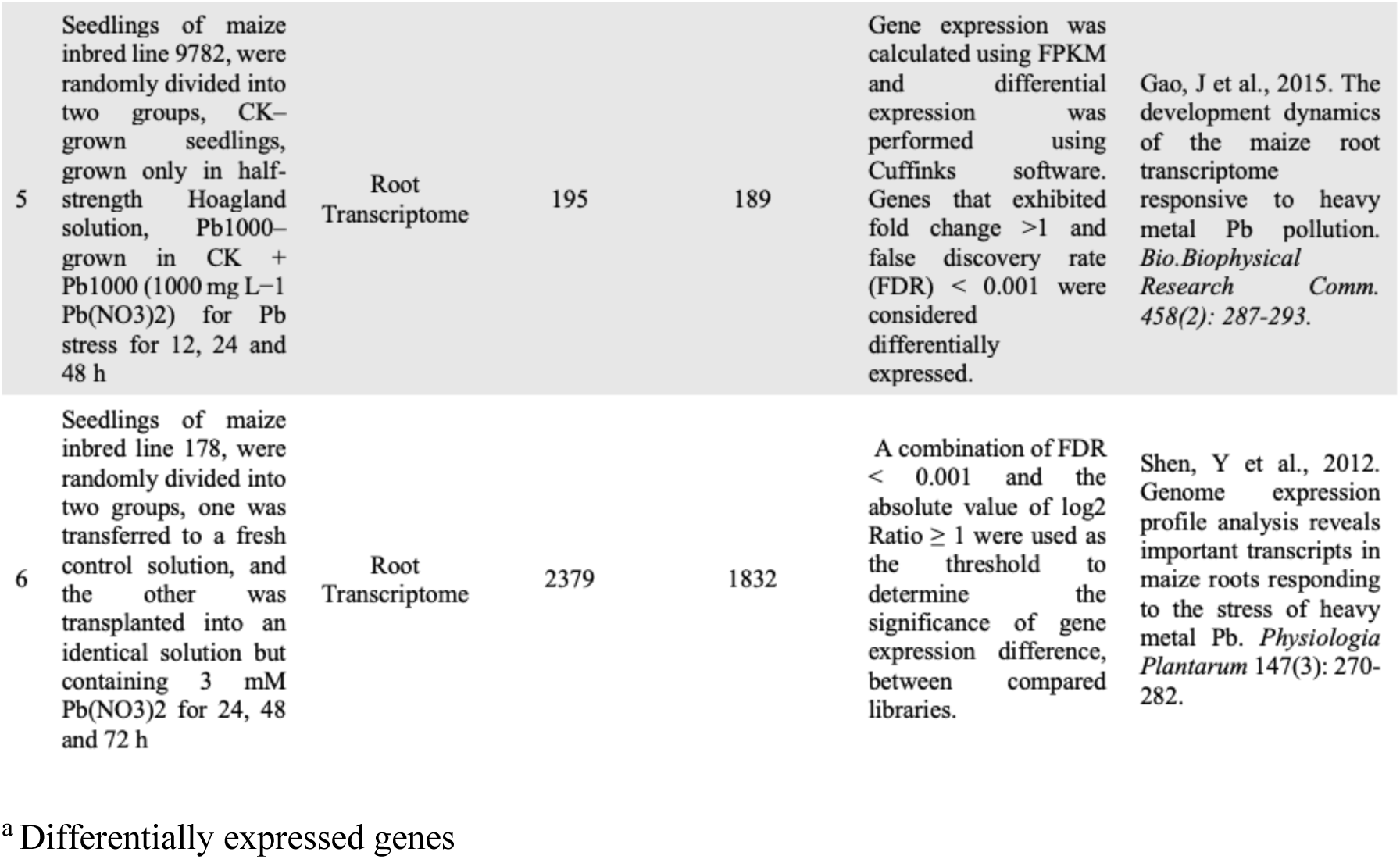
Description of six transcriptomes reporting HM response genes under different growing conditions and developmental stages.

**Table S6.**
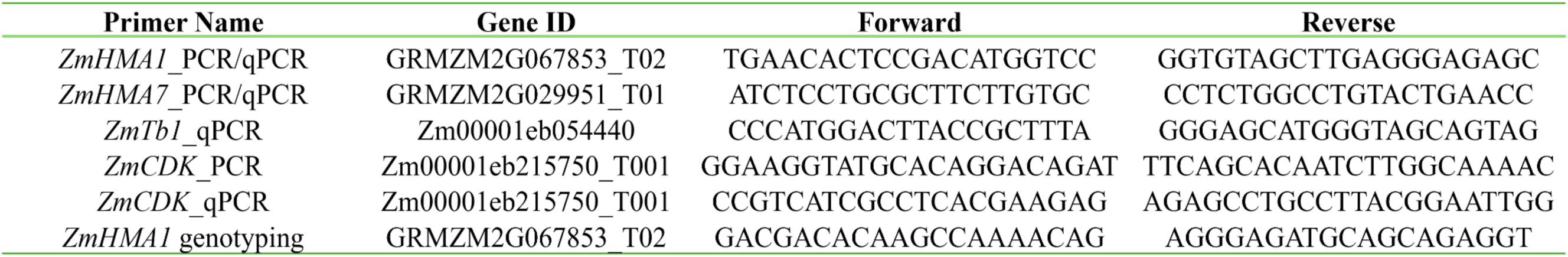
Collection of primers used in this study.

**Table S7.** Maize landrace accessions used in the genetic diversity analysis chr.5.

|  |  |
| --- | --- |
| 1 | BKN009 |
| 2 | BKN010 |
| 3 | BKN011 |
| 4 | BKN014 |
| 5 | BKN015 |
| 6 | BKN016 |
| 7 | BKN017 |
| 8 | BKN018 |
| 9 | BKN019 |
| 10 | BKN020 |
| 11 | BKN022 |
| 12 | BKN023 |
| 13 | BKN025 |
| 14 | BKN026 |
| 15 | BKN027 |
| 16 | BKN029 |
| 17 | BKN030 |
| 18 | BKN031 |
| 19 | BKN032 |
| 20 | BKN033 |
| 21 | BKN034 |
| 22 | BKN035 |
| 23 | BKN040 |

**Table S8.** Teosinte *parviglumis* accessions used in the genetic diversity analysis chr.5.

|  |  |
| --- | --- |
| 1 | TIL01 |
| 2 | TIL01-JD |
| 3 | TIL02 |
| 4 | TIL03 |
| 5 | TIL04-TIP285 |
| 6 | TIL04-TIP454 |
| 7 | TIL05 |
| 8 | TIL06-TIP260 |
| 9 | TIL06-TIP496 |
| 10 | TIL07 |
| 11 | TIL09 |
| 12 | TIL10 |
| 13 | TIL11 |
| 14 | TIL12 |
| 15 | TIL14-TIP498 |
| 16 | TIL15 |
| 17 | TIL16 |
| 18 | TIL17 |

## Notes

https://elifesciences.org/reviewed-preprints/105858#d1e1693

### Competing Interest Statement

The authors have declared no competing interest.

### Summary of Updates

Large scale genomic and transcriptomic comparison results, paleoenvironmental results and geological results have been added; Abstract updated; introductory paragraph updated; Figure 5 added; Tb1 section updated; Supplemental files 6 and 7 added

